# The SLC15A4–LAMTOR1 interaction licenses endolysosomal TLR-mediated mTOR signaling and inflammatory cytokine production

**DOI:** 10.64898/2026.06.04.729981

**Authors:** Tzu-Yuan Chiu, Peter Rory Hall, Daniel C. Lazar, Arthur S. Kim, Stefano Forli, Andrew B. Ward, John R. Teijaro, Christopher G. Parker

**Author notes:** Corresponding author: Christopher G. Parker **Email**. **Author Contributions:** C.G.P. conceived of and supervised this study. T.-Y.C. designed and performed experiments and related analysis. D.C.L. and A.S.K. contributed to reagents and assay setup. P.R.H. provided AlphaFold3 prediction, analysis, and related figures. T.-Y.C., P.R.H., S.F., A. B. W., J.R.T., and C.G.P. contributed to the preparation of the original manuscript and the preparation of the figures. All authors assisted with the review and editing of the manuscript. **Competing Interest Statement:** C.G.P., J.R.T., and D.C.L. are inventors on a patent application (US20230109595A1) submitted by The Scripps Research Institute that covers small-molecule inhibitors of SLC15A4.

## Abstract

Endolysosomal Toll-like receptors (TLRs) 7–9 are critical mediators of antimicrobial immunity, but their dysregulation drives pathogenic inflammation in autoimmune diseases such as systemic lupus erythematosus (SLE). The endolysosomal solute carrier SLC15A4 is essential for TLR7–9-mediated cytokine production and is strongly linked to SLE susceptibility, yet our understanding of the precise mechanisms by which it coordinates innate immune signaling remains incomplete. Here, we identify a direct interaction between SLC15A4 and LAMTOR1, a core scaffold subunit of the lysosomal Ragulator complex, and show that is required for coupling endolysosomal TLR activation to mTORC1 and downstream inflammatory responses. Structural modeling and site-directed mutagenesis suggest that the α1 helix of LAMTOR1 engages the substrate-binding pocket of SLC15A4, and disruption of this interface broadly suppresses TLR7–9-mediated cytokine production. Notably, this interaction regulates mTOR activation, IRF5/7 signaling, and downstream transcriptional responses through stimulus-dependent mechanisms. These findings define a previously uncharacterized SLC15A4–LAMTOR1 signaling interface that coordinates endolysosomal TLR–mTOR coupling and shapes downstream cytokine output.

**Significance Statement:** Dysregulated activation of endolysosomal Toll-like receptors (TLRs) 7–9 drives pathogenic inflammatory cytokine responses that underlie autoimmune diseases such as systemic lupus erythematosus. Despite the established role of the endolysosomal transporter SLC15A4 in this process, how it coordinates downstream immune signaling has remained incompletely understood. We show that SLC15A4 directly interacts with LAMTOR1, a core component of the lysosomal Ragulator complex, and that this interaction is required to activate mTOR and engage IRF5/7-dependent inflammatory programs downstream of TLR7–9. Disrupting this interface suppresses type I interferon and cytokine production, identifying the SLC15A4–LAMTOR1 interaction as a central node in endolysosomal innate immune signaling and validating it as a compelling target for therapeutic intervention in interferon-driven autoimmune disease.

## Introduction

The innate immune system relies on germline-encoded pattern-recognition receptors to detect microbial pathogens and initiate protective inflammatory responses. Endolysosomal Toll-like receptors (TLRs) 7, 8, and 9 sense microbial RNA and DNA and induce the production of type I interferons (IFNs) and inflammatory cytokines that restrict infection (1-4). Although this response is vital for initiating cytokine and IFN programs against pathogens, misrecognition of self-derived nucleic acids can trigger chronic inflammation and promote autoimmune and autoinflammatory diseases (5-7). Failures in immune tolerance allow self-reactive lymphocytes to survive, and aberrant TLR signaling can further amplify inflammatory pathways, particularly through endolysosomal TLR7–9 induced type I IFN production, and the induction of autoantibodies by B cells (8-10). Sustained type I IFN activity is a defining feature of several autoimmune diseases, such as systemic lupus erythematosus (SLE) and Sjögren’s syndrome (11-13). Therefore, understanding the molecular mechanisms that govern TLR7–9 signaling is essential for defining the biological basis of interferon-driven autoimmunity as well as for identifying new therapeutic targets.

SLC15A4 is a 12-transmembrane protein predominantly expressed in antigen-presenting cells, such as plasmacytoid dendritic cells (pDCs), B cells, and monocytes (14, 15). It is a member of the SLC15 protein family, which includes the proton/histidine transporter SLC15A3 and the di-/tripeptide transporters SLC15A1 and SLC15A2 (16). SLC15A3 and SLC15A4 are localized to endolysosomes via acidic dileucine motifs and are reported to transport short peptides into the cytosol by harnessing the acidic luminal environment (16, 17). Although the full substrate profile remains undefined, SLC15A4 has also been shown to transport bacterial peptidoglycans such as MDP and Tri-DAP, and loss of *Slc15a4* has been linked to reduced NOD1-driven inflammation in colitis models (16, 18-21). SLC15A4 also plays a critical role in endolysosomal TLR signaling (14, 15, 19, 22, 23). Mice deficient in *Slc15a4* exhibit normal immune cell development but display profound defects in TLR7/9-mediated production of type I IFNs, TNFα, IL-6, and IL-12 (14, 19, 24). This decreased cytokine production is not due to impaired ligand uptake or IFN secretion but stems from disruption of downstream NF-κB, IRF5 and IRF7 pathways (14, 15, 22, 23). Importantly, *Slc15a4*-deficient mice show marked protection in both the pristane-induced and spontaneous lupus models and have reduced disease and prolonged survival (15, 24-26). Despite the sequence similarity of SLC15A4 with SLC15A3 (∼60%), only the loss of SLC15A4 confers protection in these models, suggesting distinct functional roles or non-redundant pathways (15). Supporting its relevance in human disease, genome-wide association studies (GWAS) have linked SLC15A4, but not SLC15A3, to inflammatory diseases, including SLE and intestinal bowel diseases (27-31).

Despite these established roles, the mechanisms by which SLC15A4 coordinates innate immune signaling remain poorly defined and are likely multifaceted (15, 19, 22, 23, 26, 32-34). Multiple studies have linked the role of SLC15A4-mediated transport on TLR7–9 signaling by regulating endolysosomal homeostasis and mTORC1 activity (15, 19, 23, 32, 33). It has been proposed that SLC15A4 deficiency in B cells reduces lysosomal acidity and compromises vacuolar H^+^-ATPase (v-ATPase) activity, a key regulator of endocytic acidification and mTOR activity, and disrupts subsequent IRF7 activation (15, 23, 32, 35). SLC15A4 also mediates the trafficking of nucleic acid–sensing TLRs to endolysosomes, leading to defects in receptor engagement and efficient signaling (15). In addition to transport functions, SLC15A4 binds TASL, a lysosomal adaptor that recruits IRF5 and promotes its phosphorylation and downstream cytokine production (22, 26, 34). More recently, we and others have shown that SLC15A4 associates with Ragulator proteins, possibly by facilitating mTOR activation in immune cells (32, 35). Notably, we developed small molecules that appear to act by directly targeting SLC15A4 and blocking interactions with Ragulator proteins, impairing IRF5/7 activation without compromising lysosomal acidity, and thereby implicating Ragulator association to be critical for SLC15A4-mediated signaling (35, 36). Together, these findings suggest a model in which SLC15A4 functions not only as a transporter but also as a scaffold that coordinates key distinct signaling hubs to drive TLR-dependent IRF activation. While the SLC15A4–TASL–IRF5 axis has been well characterized in recent years (22, 34, 37), how SLC15A4 mediates mTOR activation warrants further studies as aberrant mTOR-dependent IFN signaling drives autoimmune pathologies (38-41).

Here, we investigate how SLC15A4 regulates mTOR activation in response to endosomal TLR stimulation. We demonstrate that SLC15A4 associates with members of the Ragulator complex, specifically LAMTOR1. Disruption of the SLC15A4-LAMTOR1 interaction with point mutations impairs mTORC1 activity and IRF5/7 responses, leading to reduced inflammatory cytokine production. Collectively, these findings define an SLC15A4–LAMTOR1 signaling interface that couples endolysosomal TLR activation to mTORC1 and shapes downstream IRF5/7-dependent cytokine output.

## Results

### SLC15A4 mediates endolysosomal TLR-induced mTOR activation and cytokine production through stimulus-dependent mechanisms

mTOR is critical for TLR-mediated expression of type I IFNs and other pro-inflammatory cytokines in various immune cells (38, 39). Numerous studies have shown that the loss of SLC15A4 disrupts mTOR signaling and impairs IRF7-dependent IFN responses (23, 32, 33). To understand how SLC15A4 contributes to endosomal TLR-mediated mTOR activation, we first assessed cytokine production after TLR7 and TLR9 activation in pDC-like CAL-1 cells, using the well-characterized ligands R848 and CpG-A/B, respectively. Consistent with prior genetic studies, IFNβ and IL-6 production in response to both TLR7/8 and TLR9 agonists was significantly reduced in SLC15A4 knockout (KO) cells (**Fig. 1A, Fig. S1, S2A-E**) (22, 34). To characterize how SLC15A4 impacts mTOR activity, SLC15A4 KO cells were treated with R848 or CpG-B. Interestingly, we observed that the phosphorylation of the mTORC1 substrate S6K remained largely unchanged (**Fig. S2F, G**). In contrast, CpG-A stimulation in SLC15A4 KO cells resulted in markedly reduced S6K phosphorylation, and diminished STAT1 phosphorylation (**Fig. 1B, C**), suggesting that CpG-A–induced mTOR activation is SLC15A4-dependent. Critically, mTOR and STAT1 phosphorylation in response to the TLR3 ligand poly(I:C) were unaffected in SLC15A4 KO cells, suggesting that SLC15A4 is specifically required for TLR9 mediated mTOR activation (**Fig. 1D, E**).

**Figure 1.**
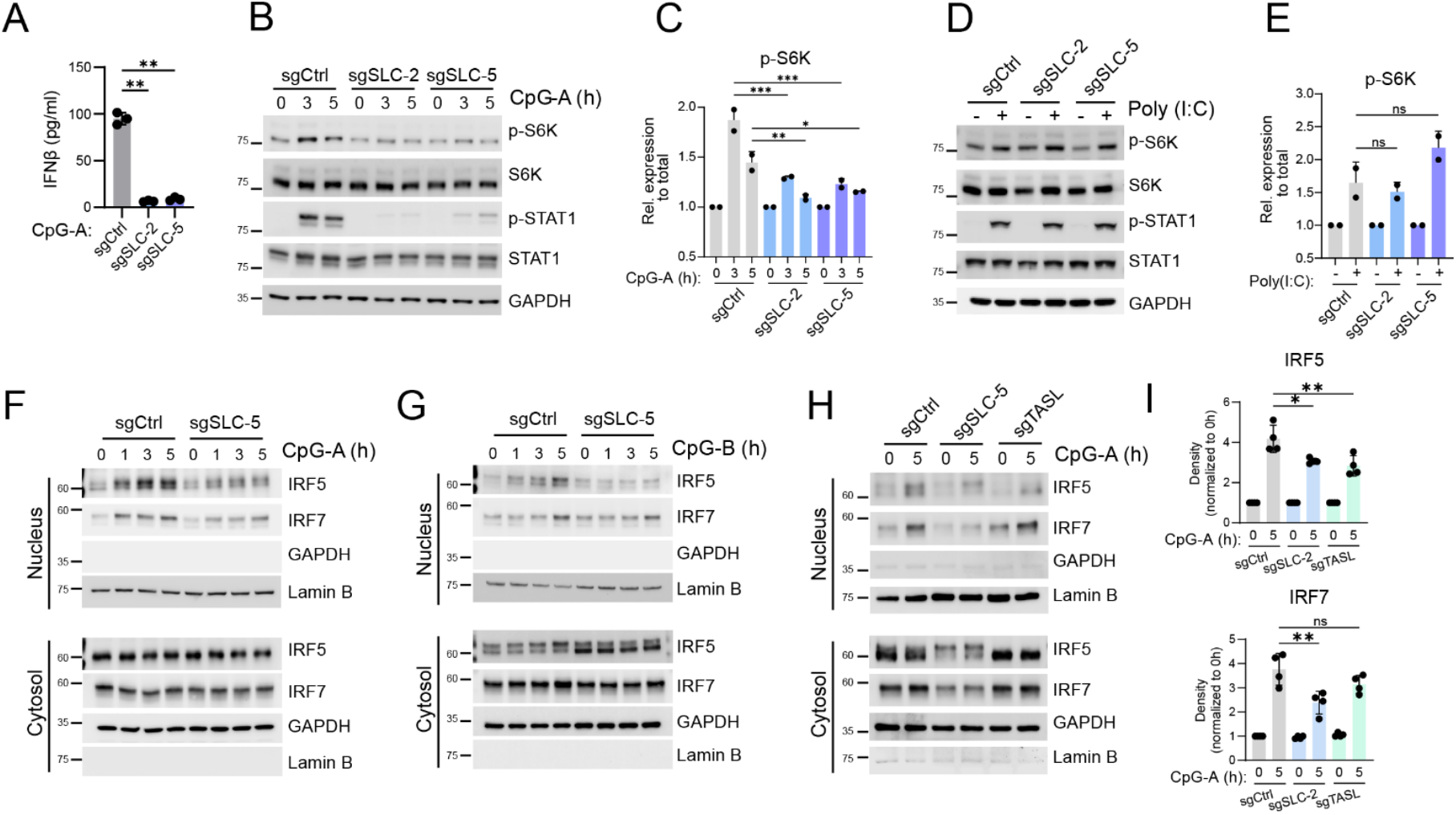
SLC15A4 mediates endolysosomal TLR-induced mTOR and cytokine production via multiple circuits. (A) IFNβ production of indicated CAL-1 cells following stimulation with CpG-A for 24 h. Data are presented as mean ± s.d. from n = 3 biological replicates. sgCtrl, single-guide RNA targeting Renilla; sgSLC-2 and sgSLC-5, two independent SLC15A4-knockout clones. (B-C) Immunoblot analysis and quantification of band intensities from the indicated CAL-1 cells stimulated with CpG-A for 0–5 h. p-, phosphorylated. (D-E) Immunoblot analysis and quantification from the indicated CAL-1 cells stimulated with poly(I:C) for 3 h. p-, phosphorylated. (F-G) Nuclear fractionation from the indicated CAL-1 cells following stimulation with (F) CpG-A and (G) CpG-B. GAPDH, cytosolic fraction control; Lamin B, nuclear fraction control. Quantification of nuclear IRFs is shown in Fig. S3A. (H-I) Nuclear fractionation and quantification of band intensities from control, SLC15A4-knockout (clone 2) and TASL-knockout CAL-1 cells stimulated with CpG-A for 5h. Statistical comparisons were performed using a one-way ANOVA test. *, p ≤ 0.05; **, p ≤ 0.01, ***, p ≤ 0.001; ns, not significant.

Both CpG-A and CpG-B activate TLR9 but trigger distinct cellular responses due to differences in endosomal localization and signaling cascade recruitment (42, 43). In primary pDCs, CpG-A accumulates in early endosomes or LC3-positive phagosomes, where it activates TLR9 and preferentially recruits IRF7, leading to robust type I IFN production. In contrast, CpG-B traffics to late endosomes and induces weaker type I IFN responses, while promoting strong proinflammatory cytokine production that is largely driven by IRF5 and NF-κB (43-47). Given that IRF5 and IRF7 are key transcriptional drivers of TLR-induced cytokine production, we next assessed the impact of SLC15A4 on their activation by monitoring nuclear translocation upon CpG-A or CpG-B stimulation in CAL-1 cells. CpG-A stimulation induced robust nuclear translocation of both IRF5 and IRF7 at 1, 3, and 5 h (**Fig. 1F, Fig. S3A**) (23, 34). In contrast, CpG-B stimulation predominantly promoted IRF5 nuclear localization, while nuclear IRF7 was only detectable at 5 h post-stimulation (**Fig. 1G, Fig. S3B**). Despite these differences, both nuclear IRF5 and IRF7 were reduced in SLC15A4-KO cells in response to both CpG-A and CpG-B (**Fig. 1F, G**). These observations suggest that CpG-A and CpG-B may have disparate impact on IRF5 and IRF7 activation.

Recent studies have identified TASL as an adaptor that links SLC15A4 to IRF5 activation by recruiting IRF5 to endolysosomes and promoting its phosphorylation (22) (37). However, our observation that *SLC15A4* loss impairs both IRF5 and IRF7 activation raises the question of whether additional SLC15A4-dependent mechanisms contribute to IRF7 activation independently of TASL. To test this, we generated TASL-deficient CAL-1 cells (**Fig. S3C)**. In agreement with previous reports, deletion of TASL abolished R848 and CpG-B-induced IRF5 nuclear translocation (**Fig. S3D-G**). Notably, *TASL* loss appears to impair CpG-B-induced IRF7 translocation, but had little effect on IRF7 nuclear translocation upon CpG-A stimulation (**Fig. 1H, I**). Together, these data suggest that while TASL primarily drives IRF5 activation across stimuli, its contribution to IRF7 is stimulus-dependent — dispensable for CpG-A but required for CpG-B-induced IRF7 activation — indicating that SLC15A4 likely supports IRF7 through additional stimulus-specific mechanisms.

### SLC15A4 associates with components of the Ragulator complex

To further investigate the potential mechanism underlying TASL-independent SLC15A4 signaling, we set out to identify additional potential binding partners of SLC15A4. Previous studies using proximity labeling and overexpression-based systems have suggested that SLC15A4 associates with components of the Ragulator-Rag complex (35, 36). To profile endogenous SLC15A4 interaction partners, we generated a polyclonal antibody targeting the N-terminus of endogenous SLC15A4 and performed immunoprecipitation (IP) followed by TMT-based mass spectrometry (MS) in CAL-1 cells (**Fig. 2A, Dataset 1**). We identified 512 enriched proteins (>5-fold over an isotype control; **Fig. 2B**), including SLC15A4 and multiple endolysosomal signaling proteins. Notably, the Ragulator subunits (LAMTOR1, -3, and -4) were significantly enriched.

**Figure 2.**
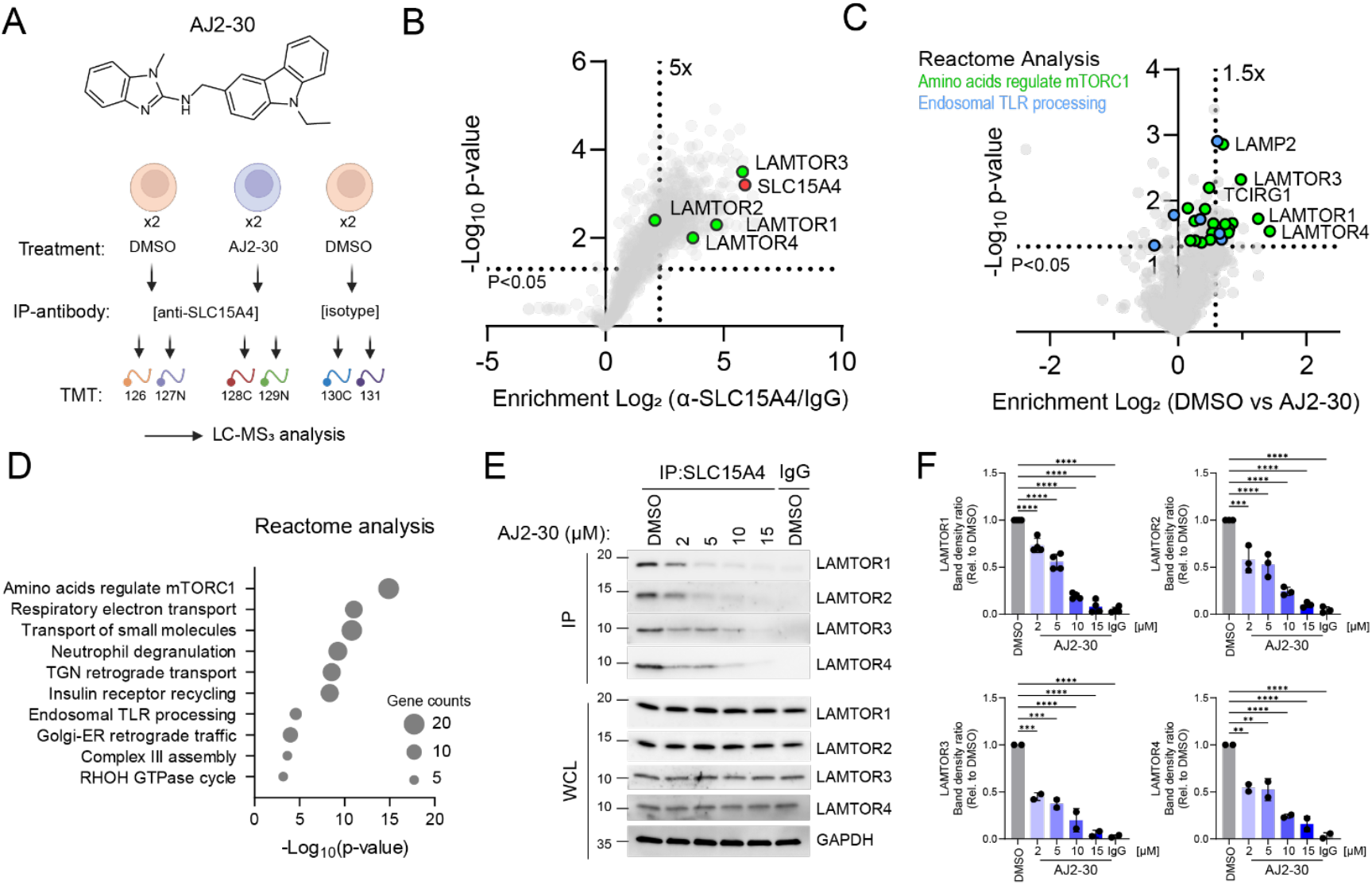
SLC15A4 interacts with components of the Ragulator complex. (A) Schematic of IP-MS experimental workflow. CAL-1 cells were treated with DMSO or AJ2-30 (5 μM) for 1h. Cell lysates were immunoprecipitated with an anti-SLC15A4 antibody and subjected to enrichment proteomics as described in Materials and Methods. The chemical structure of AJ2-30 is shown. (B-C) Volcano plots of IP–MS results. The x-axis shows protein enrichment for (B) anti-SLC15A4 versus isotype control and (C) DMSO versus AJ2-30 treatment, and the y-axis shows p-values from two biological replicates. Dotted lines indicate threshold for proteins to be designated. In B, LAMTOR1-4 are colored green, and SLC15A4 (bait) is colored red. In Fig. 2C, Amino acid regulates mTORC1 (from REACTOME analysis from Fig. 2D) proteins are shown in light green. Endosomal TLR processing proteins are shown in blue. All other proteins are plotted as gray dots. Data are presented as mean of replicated biological experiments (n = 2, P < 0.05, unpaired two-tailed t-test). (D) Reactome pathway analysis of AJ2-30 competed interactions. Pathways meeting the significance threshold (P < 0.05) are ranked by statistical significance (x axis, −log_10_ p-value). Bubble size denotes the number of proteins mapped to each pathway. Enriched pathways highlight mTORC1 regulation, vesicular trafficking, and innate immune signaling. (E-F) Immunoprecipitation of endogenous SLC15A4 in CAL-1 cells followed by immunoblot analysis. WCLs, whole cell lysates; IP, immunoprecipitation. Quantification of IP western blots are shown in Fig. 2F. Results are representative of at least two independent experiments. Statistical comparisons were performed using a one-way ANOVA test. **, p ≤ 0.01, ***, p ≤ 0.001; ****, p ≤ 0.0001.

To further prioritize functionally relevant interactions, we examined the effect of the SLC15A4-targeting compound AJ2-30 (**Fig. 2A**) (35). Given that AJ2-30 disrupts SLC15A4-dependent signaling, we reasoned that any interactions blocked by AJ2-30 would be relevant to SLC15A4 activity. Among SLC15A4-enriched proteins, 27 proteins showed significant displacement following AJ2-30 treatment (>1.5-fold decrease over DMSO), with Ragulator components LAMTOR1–4 among the most strongly disrupted interactions (**Fig. 2C, Fig. S4B**). Reactome pathway analysis of the SLC15A4 interactome upon AJ2-30 competition further highlighted amino acid regulated mTORC1 signaling as the most significantly affected pathway (**Fig. 2D)**. Immune regulators such as TASL and TLR7, as well as the mTORC1-associated proteins RRAGC and TCIRG1, were also enriched, however treatment with AJ2-30 did not affect their interaction with SLC15A4 (**Fig. S4C, D**). Immunoprecipitation followed by immunoblotting confirmed SLC15A4 interacts with Ragulator components LAMTOR1-4, and that these interactions are disrupted by AJ2-30 in a dose-dependent manner (**Fig. 2E, F**). Finally, confocal imaging revealed reduced colocalization of LAMTOR1 with recombinantly expressed HA-tagged SLC15A4 upon AJ2-30 treatment, consistent with the reduced SLC15A4-LAMTOR1 interaction observed by IP experiments (**Fig. S4E, F**). Together, these data demonstrate that SLC15A4 associates with Ragulator components in a manner sensitive to pharmacological disruption of SLC15A4 function, as well as other immune regulators.

### LAMTOR1 links TLR7-9 to the mTOR–IRF5/7 signaling axis

Ragulator is a pentameric complex composed of LAMTOR1–5, in which LAMTOR1 wraps around LAMTOR2-5 and serves as a scaffold to anchor the entire complex to the lysosomal membrane, thus enabling Rag GTPase recruitment and mTORC1 activation (48-53). Given the critical role of LAMTOR1 in this regard, as well as the established roles of mTORC1 signaling for TLR-induced cytokine production (38-41), we next investigated whether the interaction between SLC15A4 and LAMTOR1 was required for TLR7-9 mediated responses. We first generated LAMTOR1-KO CAL-1 cells using CRISPR–Cas9 gene-editing (**Fig. 3A**). Loss of LAMTOR1 led to a substantial reduction in LAMTOR2–5 protein levels, whereas other endolysosomal protein levels, including SLC15A4, remained unchanged (**Fig. S5A, B**) (54, 55). LAMTOR1-deficient cells displayed significantly reduced IFNβ secretion upon TLR9 stimulation with CpG-A and CpG-B (**Fig. 3B, C**) as well as reduced IFNβ and TNFα secretion upon R848 stimulation (**Fig. 3D**). We noted that depletion of LAMTOR1 abrogated phosphorylation of mTORC1 substrates S6K under both basal and stimulated conditions, however, only impaired STAT1 phosphorylation upon TLR9 stimulation but not TLR7, pointing to stimulus-dependent mechanisms downstream of LAMTOR1 **(Fig. S6A-G)**. Importantly, nuclear translocation of IRF5 and IRF7 was abrogated upon CpG-A and CpG-B stimulation in LAMTOR1-KO cells (**Fig. S7A-D**) but was unaffected following R848 stimulation **(Fig. S7E, F)**. Consistent with these findings, qPCR analysis showed that R848-induced *TNFA* and *IFNB1* mRNA was not impaired, despite reduced cytokine secretion at early time points (0–7 h) (**Fig. S7G, H)**. Collectively, these data point to a critical role of LAMTOR1 in endolysosomal TLR-mediated cytokine production, albeit through distinct mechanisms depending on the stimulus. While the impact of LAMTOR1 on TLR9-mediated cytokine production occurs via a mTOR-IRF5/7 signaling axis, its effect on TLR7-mediated responses likely operates at a post-transcriptional level, consistent with established roles of mTOR in translational and secretory regulation (38, 56).

**Figure 3.**
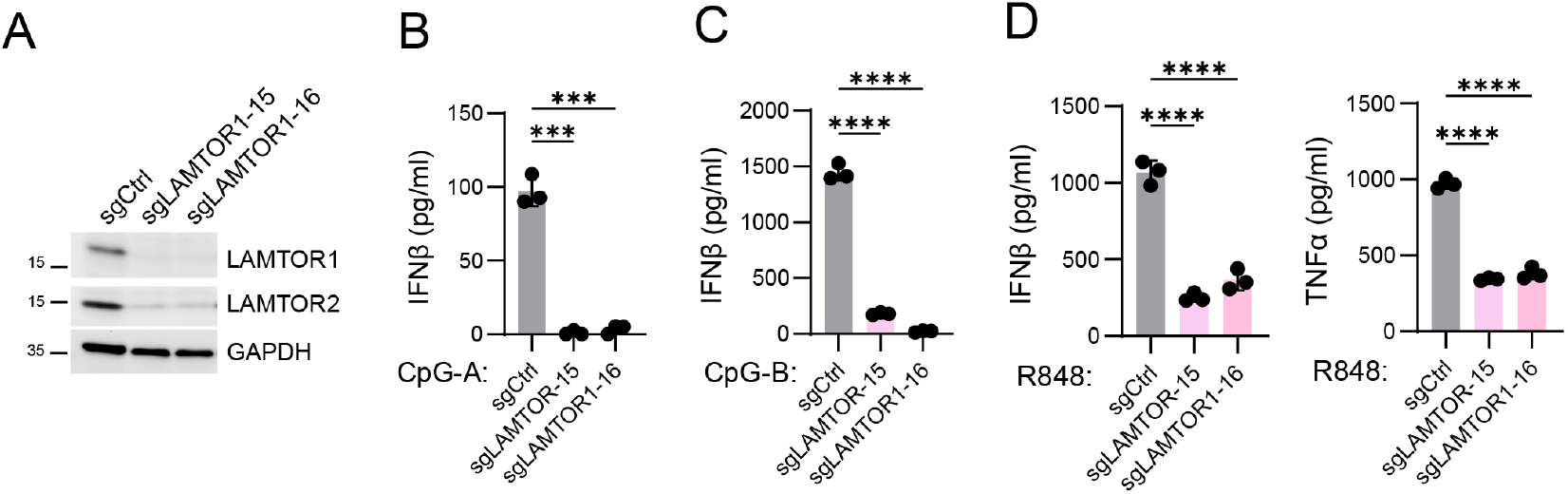
LAMTOR1 is required for TLR7-9-induced cytokine production. (A) Immunoblot analysis and quantification from the indicated CAL-1 cells. sgCtrl, single-guide RNA targeting Renilla; sgLAMTOR1-15 and sgLAMTOR1-16, two independent LAMTOR1-knockout clones. FL, full length. (B-D) Cytokine production in the indicated CAL-1 cells following stimulation with (B) CpG-A, (C) CpG-B, and (D) R848 for 24 h. Data are presented as mean ± s.d. from n = 3 biological replicates. Statistical comparisons were performed using a one-way ANOVA test. ***, p ≤ 0.001; ****, p ≤ 0.0001.

### The LAMTOR1-SLC15A4 interface mediates TLR9–mTOR signaling and cytokine production

To further resolve potential SLC15A4-mediated effects on TLR-induced mTOR activity, we next sought to identify the key residues required for the SLC15A4 and LAMTOR1 interaction. To probe this interaction, we first demonstrated that LAMTOR1-deficient cells reconstituted with Myc-tagged wild-type (WT) LAMTOR1 led to restoration of TLR9 responses (**Fig. S8A**) and mutation of LAMTOR lipidation residues Cys3/4 (CCSS), which are required for LAMTOR1 lysosomal anchoring, markedly reduced LAMTOR1 protein stability and abolished its ability to restore CpG-A induced IFNβ production (**Fig. 4C; Fig. S8F**). Next, to understand the specific structural determinants of the SLC15A4-LAMTOR1 interaction, we generated multiple models with different seeds using AlphaFold3 (57), in which seed three predicted insertions of the LAMTOR1 α1-helix into the SLC15A4 substrate pocket (**Fig. 4A; Fig. S9)**, analogous to the α-helix of the N-terminus of TASL (58). Based on this prediction, we generated truncations removing α1 (amino acids 40–50 [Δ40-50], 40–65 [Δ40-65], 40–80 [Δ40-80], or 40–95 [Δ40-95]) or α3/α4 (Δ95–161) (**Fig. 4B)**. Deletion of the α1 helix region, but not those in α3/α4, abolished SLC15A4 binding to LAMTOR1 without affecting protein stability (**Fig. 4C; Fig. S8B-E**). Moreover, the α1-truncated mutants failed to rescue IFN-β production upon CpG-A stimulation (**Fig. S8F**) but also displayed reduced basal mTOR signaling (**Fig. S8G-H**).

**Figure 4.**
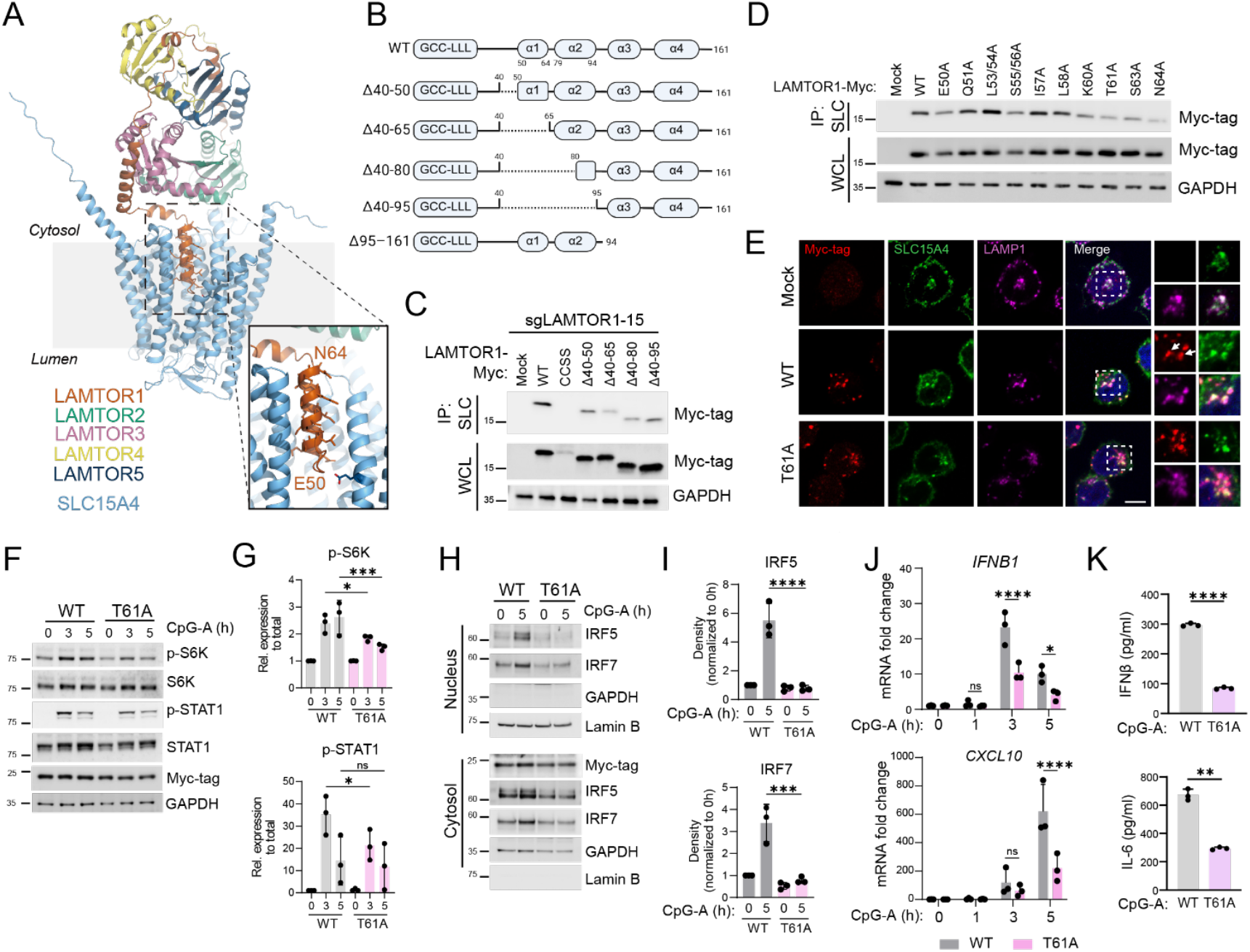
LAMTOR1 α1-helix interacts with SLC15A4 and is required for endolysosomal TLR mediated signaling. (A)Representative AlphaFold3 model of the complex between SLC15A4 and the Ragulator (LAMTOR1–5). Potential binding interfaces are zoomed in and LAMTOR1 α1-helix residues E50 and N64 are highlighted in brown. (B)Schematic representation of LAMTOR1 middle-region deletion mutants (Δ40–50, Δ40–65, Δ40–80, Δ40–95, and Δ95–161). (C)Immunoprecipitation of endogenous SLC15A4 from LAMTOR1-knockout CAL-1 cells reconstituted with the indicated WT and truncated Myc-LAMTOR1 constructs, followed by immunoblot analysis. Mock denotes vector control. CCSS denotes mutations of Cys3 and Cys4 in LAMTOR1. (D)Immunoprecipitation of endogenous SLC15A4 from LAMTOR1-knockout CAL-1 cells reconstituted with the indicated WT and mutant Myc-LAMTOR1 constructs, followed by immunoblot analysis. (E) Immunostaining of endogenous SLC15A4 and reconstituted-LAMTOR1 (Myc-tag) in indicated CAL-1 cells. Cells were co-stained with LAMP1 as a lysosomal marker. Zoom-in regions are indicated by white dashed boxes. Colocalized regions of SLC15A4 and Myc-tag are indicated by white arrows. Scale bar, 5 μm. (F-G) Immunoblots and quantification of LAMTOR1-knockout CAL-1 cells reconstituted with the indicated WT and mutant Myc-LAMTOR1 constructs upon stimulation with CpG-A for 0–5 h. p-, phosphorylated. (H-I) Nuclear fractionation and quantification of LAMTOR1-knockout CAL-1 cells reconstituted with the indicated WT and mutant Myc-LAMTOR1 constructs upon stimulation with CpG-A for 0–5 h. (J)*IFNB1* and *CXCL10* mRNA levels in the indicated CAL-1 cells following CpG-A stimulation, measured by RT–PCR. Fold changes were calculated relative to the unstimulated group. (K)Cytokine production of indicated CAL-1 cells stimulated with CpG-A for 24 h. Data are presented as mean ± s.d. from at least two biological replicates. Statistical comparisons were performed using a two-tailed Student’s t-test. *, p ≤ 0.05; **, p ≤ 0.01; ***, p ≤ 0.001; ****, p ≤ 0.0001; ns, not significant.

To further refine this interaction, we performed alanine mutagenesis of the α1 helix and reconstituted these mutants in LAMTOR1-deficient cells. We identified a cluster of residues spanning Lys60-Asn64 on LAMTOR1 that impeded interactions with endogenous SLC15A4 without impacting SLC15A4 or TLR9 expression levels (**Fig. 4D; Fig. S10A-C**). Notably, these mutants reduced levels of S6K phosphorylation upon CpG-A stimulation but had minimal impact on basal levels (**Fig. S10E-F**). Among these mutants, T61A showed among the greatest reduction in SLC15A4 binding, accompanied by reduced colocalization with SLC15A4 despite preserved lysosomal localization (**Fig. 4E; Fig. S10D**). Consistent with impaired SLC15A4 engagement, T61A cells displayed reduced S6K and STAT1 phosphorylation upon CpG-A stimulation but not under basal conditions, corroborating our observations in SLC15A4-deficient cells (**Fig. 4F-G; Fig. S10E-F**). In contrast to CpG-A, we observed no significant impact on mTOR or STAT1 signaling upon R848 or CpG-B stimulation in T61A cells (**Fig. S11A-D**) and no disruption of mTOR signaling was observed upon stimulation of poly(I:C) (**Fig. S10G, H**), suggesting these effects are largely selective for CpG-A stimulus. At the level of IRF activation, the T61A mutant failed to restore CpG-A–induced IRF5/7 nuclear translocation, with more modest effects following CpG-B and no apparent defects following R848 stimulation (**Fig. 4H-I, Fig. S11E-H**). Consistently, induction of IRF5/7 target genes *IFNB1* and *CXCL10* was significantly reduced upon CpG-A stimulation, while CpG-B–induced *IFNB1* showed less pronounced reduction and R848-induced *IFNB1* was modestly increased (**Fig. 4J, Fig. S11I-J**) (59). In line with transcriptional results, production of cytokines, including IFNβ, IL-6, and TNFα was substantially reduced following CpG-A stimulation while less pronounced reduction was observed in response to CpG-B and R848 (**Fig. 4K, Fig. S12A, B**). Interferon-induced responses to TLR3 or IFNAR stimulation remained intact (**Fig. S12C, D)**. Together, these data identify the LAMTOR1 α1 helix as a structurally and functionally essential determinant of TLR7–9-induced cytokine production, with effects most prominent for CpG-A-driven responses.

Previous studies showed that the E465K mutation in SLC15A4 prevents interaction with TASL (22, 23, 60, 61). To determine whether this interface is also required for LAMTOR1 engagement, we reintroduced HA-tagged wild-type SLC15A4 or the E465K mutant into SLC15A4-deficient CAL-1 cells. While co-immunoprecipitation of SLC15A4 resulted in strong co-enrichment of LAMTOR1, the E465K mutant exhibited a significant reduction in binding to LAMTOR1 (**Fig. 5A; Fig. S13A, B**). Importantly, this impaired LAMTOR1 interaction with SLC15A4 was not dependent on TASL engagement, as the interaction between SLC15A4 and LAMTOR1 was maintained in TASL-knockout cells (**Fig. S13C)**. Consistent with these results, we observed reduced co-localization of SLC15A4 E465K with LAMTOR1 relative to wild-type SLC15A4 (**Fig. 5B, Fig. S13D**). When HA-tagged SLC15A3 was expressed in SLC15A4-deficient cells, LAMTOR1 was not enriched, indicating that these observed interactions were specific to SLC15A4 **(Fig. 5A)**.

**Figure 5.**
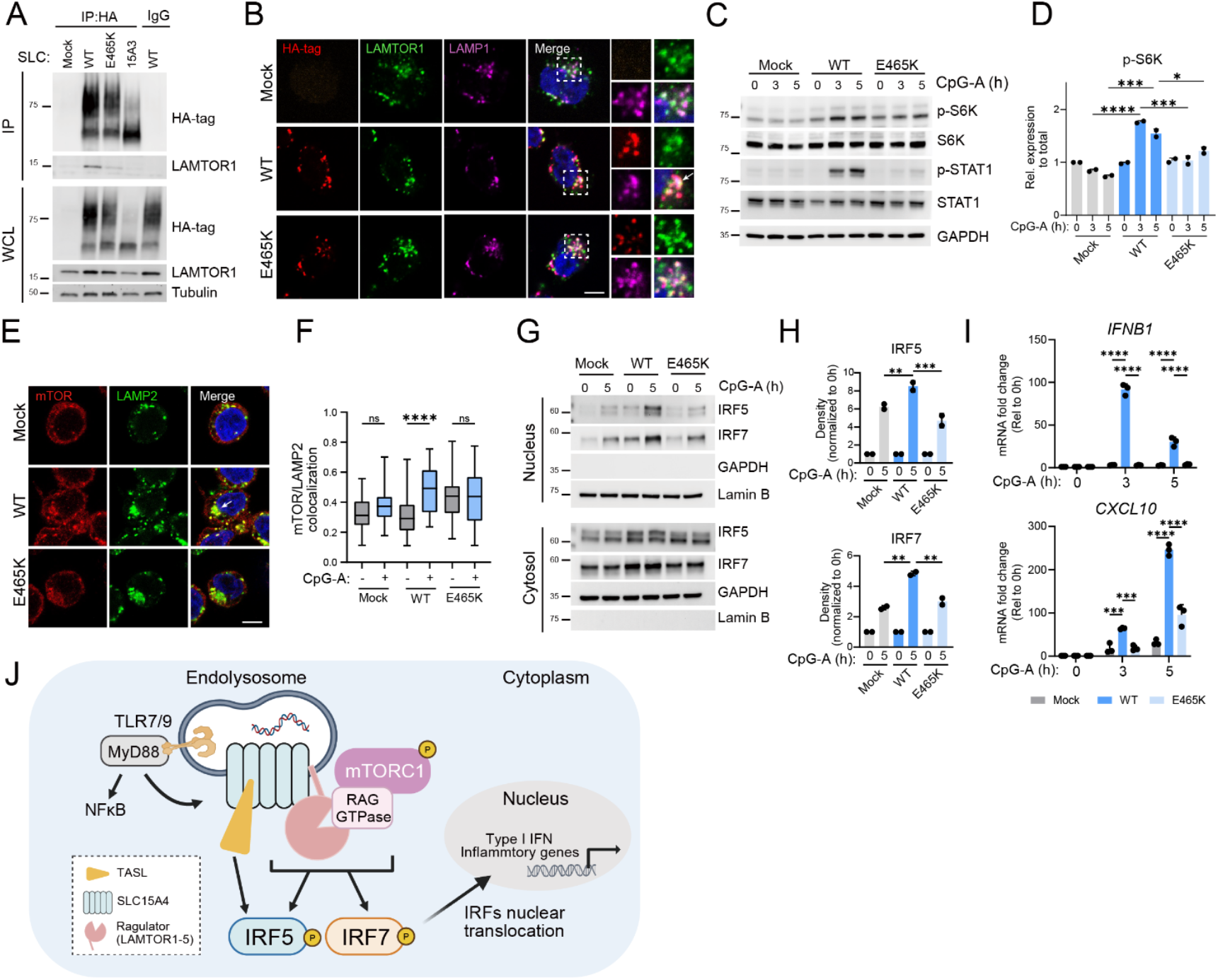
The SLC15A4 substrate-binding pocket mediates LAMTOR1 engagement. (A)Immunoprecipitation of HA-tag proteins from SLC15A4-knockout CAL-1 cells reconstituted with the indicated WT and mutant HA-tagged constructs upon stimulation, followed by immunoblot analysis. Mock denotes vector control. E465K denotes mutations of Glu465 in SLC15A4.15A3 denotes SLC15A3. (B)Immunostaining of HA-tag SLC15A4 and endogenous LAMTOR1 in CAL-1 cells. Cells were co-stained with LAMP1 as a lysosomal marker. Zoom-in regions are indicated by white dashed boxes. Colocalized regions of HA-tag and LAMTOR1 are indicated by white arrows. Scale bar, 5 μm. (C, D) Immunoblots and quantification of band intensity from the indicated CAL-1 cells stimulated with CpG-A for 0–5 h. p-, phosphorylated. (E)Immunostaining of mTOR in CAL-1 cells following CpG-A stimulation for 5 h. Cells were co-stained with LAMP2 as a lysosomal marker. Colocalized regions of mTOR and LAMP2 are indicated by white arrows. Scale bar, 5 μm. (F)Quantification of mTOR colocalization with LAMP2 in Fig. 5E. (G, H) Nuclear fractionation and quantification from the indicated CAL-1 cells stimulated with CpG-A for 5 h. (I)*IFNB1* and *CXCL10* mRNA levels in the indicated CAL-1 cells following CpG-A stimulation, measured by RT–PCR. Fold changes were calculated relative to the unstimulated group. (J)Schematic model of SLC15A4- and Ragulator-dependent IRF5/7 activation downstream of TLR7–9. Statistical comparisons were performed using a two-tailed Student’s t-test for Fig. 5E and one-way ANOVA for all other analyses. *, p ≤ 0.05; **, p ≤ 0.01; ***, p ≤ 0.001; ****, p ≤ 0.0001; ns, not significant.

Consistent with the impaired LAMTOR1 interaction, we observed that CAL-1 cells with the E465K mutation in SLC15A4 had significantly reduced S6K phosphorylation upon CpG-A stimulation (**Fig. 5C, D**), which was accompanied by impaired mTOR recruitment to lysosomes (**Fig. 5E, F**) (62). Moreover, the cells expressing the SLC15A4 E465K mutant also failed to fully rescue IRF5/7 nuclear translocation upon CpG-A stimulation (**Fig. 5G, H**). This IRF7 defect was not observed upon TASL deletion, highlighting a unique role for the LAMTOR1–SLC15A4 interaction in IRF7 activation (**Fig. 1H, I**) (37). Finally, induction of *IFNB1* and *CXCL10* was significantly reduced upon CpG-A stimulation in E465K-reconstituted cells (**Fig. 5I**). Collectively, these data establish that the SLC15A4–LAMTOR1 interaction is required for endolysosomal TLR-mediated cytokine production, most prominently coupling TLR9–CpG-A activation to the mTOR–IRF5/7 axis, with more modest contributions to CpG-B and TLR7 responses (**Fig. 5J**), which appear to be primarily affected at a post-transcriptional level.

## Discussion

mTORC1 is essential for coordinating innate immune defenses downstream of endosomal TLR signaling (40, 56). Genetic and pharmacological inhibition of mTOR activity markedly impairs TLR7-9 dependent type I IFN and inflammatory cytokine production, largely through disruption of MyD88-dependent activation of IRF5 and IRF7 (38, 39, 63). Beyond direct transcriptional regulation, mTOR signaling also supports the metabolic and biosynthetic programs required for activation of pDCs and B cells in response to endosomal TLR stimulation (64, 65). Our findings characterize the interaction between SLC15A4 and the Ragulator component LAMTOR1 that resolves how SLC15A4 couples endosomal TLR activation to mTORC1, providing a molecular mechanism for the previously described role of mTOR in IRF5/7-dependent cytokine production (23, 35). Disruption of this complex attenuates mTOR signaling and impairs downstream IRF5/7-dependent cytokine responses, underscoring a functional role for the SLC15A4–LAMTOR1 platform in mediating this pathway.

Several observations support the SLC15A4–LAMTOR1 framework. Endogenous interaction proteomics revealed robust association of SLC15A4 with Ragulator components, and these interactions were selectively disrupted by the SLC15A4-targeting compound AJ2-30 (35). Structural modeling and mutagenesis further identified a functionally important interface between the LAMTOR1 α1 helix and the SLC15A4 substrate-binding region. Importantly, perturbation of this interface impaired TLR-induced mTOR activation and downstream inflammatory signaling. Together, these findings suggest that productive coupling of endosomal TLR signaling to mTOR activity requires direct engagement of the Ragulator complex by SLC15A4, rather than basal mTOR activity alone.

A notable aspect of this signaling axis is its apparent stimulus specificity. Disruption of the SLC15A4–LAMTOR1 interface most strongly impaired CpG-A-induced mTOR activation, IRF5/7 signaling, and interferon-associated transcriptional responses, whereas CpG-B and R848 responses were comparatively less affected at the level of mTOR–IRF signaling despite persistent defects in cytokine production. These observations parallel the known compartment-specific behavior of endosomal TLR ligands. CpG-A preferentially localizes to signaling compartments that support robust IRF7-dependent type I IFN responses (47, 66, 67), whereas CpG-B and R848 are more strongly associated with NF-κB- and IRF5-skewed inflammatory outputs (43, 68, 69). Our findings therefore suggest that distinct endosomal TLR ligands engage partially separable SLC15A4-dependent signaling states, potentially reflecting differences in endosomal maturation, signaling complex assembly, or metabolic requirements.

This stimulus-selective behavior may also help reconcile the relationship between the SLC15A4–LAMTOR1 axis described here and the previously established SLC15A4–TASL–IRF5 pathway (22, 26, 34, 37). TASL has emerged as a key adaptor linking SLC15A4 to IRF5 activation downstream of TLR7–9 signaling, particularly in the context of R848 and CpG-B stimulation. In contrast, our data suggest that CpG-A-induced IRF7 and mTOR responses rely more heavily on LAMTOR1-dependent signaling mechanisms. Rather than functioning through a single linear pathway, SLC15A4 may therefore coordinate multiple signaling modules whose relative contributions differ depending on ligand trafficking and endosomal context. In this framework, TASL-associated signaling may preferentially support IRF5-dominant inflammatory responses, whereas LAMTOR1/Ragulator coupling may more strongly support IRF7 programs and type I IFN production.

Our findings also raise the possibility that SLC15A4-dependent cytokine regulation extends beyond direct transcriptional control. Although disruption of the SLC15A4–LAMTOR1 interface strongly impaired CpG-A-induced transcriptional responses, cytokine production defects following CpG-B or R848 stimulation occurred despite comparatively modest effects on IRF5/7 activation. Given the established role of mTOR in coordinating translational, metabolic, and biosynthetic programs in activated immune cells (64, 65), these observations suggest that distinct SLC15A4-associated signaling complexes may regulate inflammatory outputs at multiple levels. More broadly, these findings support the idea that endolysosomal signaling is not governed by a single uniform pathway, but instead by dynamically assembled signaling networks whose composition and functional output vary depending on ligand identity and endosomal localization.

Finally, these studies further expand the emerging view that endolysosomal solute carriers can function as signaling organizers in addition to transporter roles. The ability of SLC15A4 to associate with TLR regulators, TASL, and Ragulator components suggests that it may function as a multifunctional endolysosomal scaffolding hub that integrates receptor activation with metabolic and inflammatory signaling pathways. Such organization may be particularly important in pDCs and related immune populations that require highly coordinated type I IFN responses. This framework also has potential therapeutic implications. Genetic and pharmacological studies have established SLC15A4 as a promising target in lupus and other interferon-driven autoimmune diseases (15, 24, 25, 28, 30), and our findings suggest that disruption of all SLC15A4-associated signaling interfaces may more robustly suppress pathogenic inflammatory signaling. More generally, defining how endolysosomal signaling complexes are spatially organized may reveal new opportunities to therapeutically tune innate immune responses.

## Materials and Methods

### Cell lines and stimulation with TLR ligands

Lenti-X™293T (#632180) was purchased from Takara Bio. CAL-1 cells were obtained from Shimeru Kamihira (Nagasaki University) (70). Lenti-X™293T cells were maintained in DMEM medium supplemented with 10% FBS, 1x Pen/Strep, and 2mM L-glutamine. CAL-1 cells were maintained in RPMI 1640 medium supplemented with 10% FBS, 1x Pen/Strep, and 2mM L-glutamine. All cell lines were grown at 37 °C in a humidified 5% CO_2_ atmosphere. Unless otherwise indicated, cells were plated at 1 × 10^6^ cells/mL in RPMI 1640 supplemented with 1% FBS and rested overnight (71). The following day, 1.5 × 10^6^ cells/mL cells were stimulated with 5 μg/mL R848, 5 μM CpG-A, or 1 μM CpG-B in serum-free RPMI 1640 for the indicated durations. For extended stimulations (>6 h), cells were stimulated in RPMI 1640 containing 2% FBS.

Other Materials and Methods details can be found in the SI Appendix.

## Data, Materials, and Software Availability

Uncropped, full western blot images and gels are provided in **Fig. S14**. The mass spectrometry proteomics data have been deposited to the ProteomeXchange Consortium via the PRIDE (72) partner repository with the dataset identifier PXD073184. All other results are included in the article and/or supporting information.

(For peer reviewer access details)

Log in to the PRIDE website using the following details:

Project accession: PXD073184

Token: RKnbqRHwHnDT

Alternatively, reviewer can access the dataset by logging in to the PRIDE website using the following account details:

Username

Password: 6a6BFKyLnVHH

## Acknowledgments

We are grateful to the Scripps Research Microscopy Core Facilities and Flow Cytometry Core Facilities, and Weichao Li, Ph.D. for assisting with the IP-MS experiments. This work was supported by the National Institute of Allergy and Infectious Diseases (R01 AI156268 to C.G.P. and J.R.T.) and National Institute of General Medicine (R01 R01GM069832 to S.F.). A.S.K. was supported by TL1 DK143272 and in part by Open Philanthropy and the Life Sciences Research Foundation.

