## Supplemental Information for "The SLC15A4–LAMTOR1 interaction licenses endolysosomal TLR-mediated mTOR signaling and inflammatory cytokine production"

*Corresponding author: Christopher G. Parker

**This PDF file includes:**

Supporting text

Figures S1 to S14

Tables S1 to S4

Legend for Dataset S1

SI References

**Other supporting materials for this manuscript include the following:**

Datasets S1

**Supporting Information Text**

**Materials and Methods**

All antibodies, key reagents and kits are listed in Table S4.

Generation of anti-SLC15A4 antibody

Custom rabbit anti-SLC15A4 polyclonal antibodies were generated against the N-terminal region (amino acids 1-14) of SLC15A4 (Genscript). Rabbit serum was affinity-purified using a SLC15A4 peptide-conjugated column (Scripps Research Antibody Core Facility). Antibody specificity was validated by immunofluorescence and flow cytometry using SLC15A4-knockout cells (**Fig. S1**). Suitability for immunoprecipitation was confirmed using SLC15A4-overexpressing cells (**Fig. S4A**). Notably, this antibody did not detect endogenous or overexpressed SLC15A4 by western blot. Accordingly, flow cytometry was used to confirm equivalent SLC15A4 expression levels prior to immunoprecipitation.

Plasmids and site-directed mutagenesis

pLentiCRISPRv2 (Addgene #52961) was used to generate CRISPR-Cas9 knockout cells. sgRNAs for control (targeting *Renilla luciferase*), SLC15A4 and TASL have been described previously (1), whereas sgRNAs targeting LAMTOR1 were designed using CHOPCHOP v3 (2) . Oligonucleotides were synthesized by Integrated DNA Technologies, and sgRNA sequences are provided in *Table S1*. Site-directed mutagenesis and truncation constructs were generated using Phusion™ Flash High-Fidelity PCR Master Mix (Thermo Fisher Scientific; F548L) followed by In-Fusion® cloning (Takara Bio USA; 638947). Rescued experiments were performed using cDNA constructs where the sequences were modified to be resistant to Cas9-mediated cleavage. N-terminal HA-tagged SLC15A4 constructs were cloned into a modified pLEX_305 vector (Addgene #41390), in which the puromycin resistant gene was replaced with hygromycin resistant cassette. C-terminal Myc-tagged LAMTOR1 constructs were cloned into pLJC6-EV vector (Addgene #163454). All plasmids were validated by Sanger sequencing (Azenta) or whole-plasmid sequencing (Plasmidsaurus). Oligonucleotides used for mutagenesis are provided in *Table S2*.

Generation of stable knockout or overexpression cell lines via lentiviral transduction

Lentiviral transduction was performed as previously described (3). Briefly, Lenti-X 293T cells were transfected with sgRNA- or cDNA-encoding lentiviral vectors and packaging plasmids psPAX2 and pMD2.G (Addgene #12260 and 12259) using PEI MAX (Polysciences Inc). Culture medium was replaced 24 h after transfection, and viral supernatants were collected at 48 h and 72 h post-transfection, filtered through a 0.2-µm PES membrane, and supplemented with 8 µg/mL polybrene (Sigma) to optimize transduction efficiency. Target cells were transduced by spinoculation (800 × *g* for 30 min at room temperature). Fresh medium was added 24 h later, and antibiotic selection was initiated 48 h post-infection using puromycin (2 µg/mL), blasticidin (10 µg/mL), or hygromycin (350 µg/mL), as appropriate.

For SLC15A4 and LAMTOR1 CRISPR knockout cell lines, single-cell clones were isolated by limiting dilution and expanded from 96-well U-bottom plates after approximately 20 days (SLC15A4) or 35 days (LAMTOR1) of growth. TASL, IRF5, and IRF7 CRISPR knockout cell lines were generated and used as pooled populations following 10 days of antibiotic selection. For cDNA overexpression, antibiotic-selected pooled cells were used for experiments without further subcloning.

Immunoprecipitation-mass spectrometry (IP-MS) sample preparation

AJ2-30 (CAS 2700322-79-6) was synthesized in lab as previously described (4). 20 million CAL-1 cells were incubated with 5 μM AJ2-30 or DMSO control (in duplicate) in culture medium for an hour at 37°C. Cells were harvested, washed twice with ice-cold PBS, and resuspended in ice-cold E1C lysis buffer (50 mM HEPES, 250 mM NaCl, 5 mM EDTA, 0.3% [w/v] CHAPS) supplemented with protease inhibitors. Cells were sonicated on ice twice (15 ms on, 40 ms off, 15% amplitude; total on-time 1 s), incubated on ice for 20 min, and clarified by centrifugation at 13,000 × *g* for 10 min at 4°C. Supernatants were transferred to new tubes and incubated with 10 µg anti-SLC15A4 antibody or isotype control (BioLegend #910801) overnight at 4°C rotating, followed by incubation with 75 µL Protein A/G magnetic beads (MCE #HY-K0202) for 2 h at 4°C. Beads were washed six times with 1 mL E1C buffer. Bound proteins were eluted with 130 µL of 5% SDS in 50 mM triethylammonium bicarbonate (TEAB; Thermo Fisher Scientific #90114) by incubation at 37°C for 1 h. Eluates were collected and processed for downstream proteomic analysis.

Sample Processing and Tandem Mass Tag (TMT) Labeling for Proteomics

Immunoprecipitated samples were reduced with tris(2-carboxyethyl)phosphine (TCEP; final concentration 20 mM) for 30 min at room temperature and subsequently alkylated with iodoacetamide (final concentration 40 mM) for 15 min at room temperature in the dark. Samples were acidified with 12% phosphoric acid to a final concentration of approximately 1.1% and processed using S-Trap™ Mini Columns (PROTIFI; C02-mini-40) as previously described (5). Briefly, 900 µL S-Trap buffer (90% methanol, 100 mM TEAB, pH 7.1) was added, and samples were loaded onto S-Trap™ Mini Columns. Columns were washed three times with 400 µL S-Trap buffer. Proteins trapped on the columns were digested with 1 µg trypsin/LysC (Promega; V5071) in the presence of 0.05% (v/v) ProteaseMAX™ (Promega; V2071) overnight at 37 °C. Peptides were sequentially eluted with (1) 80 µL 50 mM TEAB (pH 8.0), (2) 80 µL 0.2% formic acid, and (3) 80 µL 0.2% formic acid in 50% acetonitrile (ACN), with each elution collected by centrifugation (10,000 × *g*, 1 min, room temperature) into Protein LoBind tubes (Eppendorf). Eluted peptides were dried using a SpeedVac vacuum concentrator (Thermo Fisher Scientific) and resuspended in 100 µL of 30% ACN in 50 mM TEAB.

The digest peptides were labeled with respective 10-plex TMT (Thermo Fisher Scientific #90406) reagents for an hour, quenched with hydroxylamine (Thermo Fisher Scientific #90115), and acidified with formic acid prior to drying under vacuum centrifugation. The dried TMT-labeled samples were resuspended in 300 μl 0.1% trifluoroacetic acid (TFA, Thermo Fisher Scientific #85183) and fractionated using a Pierce High pH Reversed-Phase Fractionation Kit (Thermo Fisher Scientific # 84868). Peptides were eluted with increasing concentrations of ACN (5%, 10%, 20%, 35%, and 50%) in 0.1% triethylamine, dried, and stored at -80°C until LC–MS analysis. The first fraction was excluded from subsequent analysis. Prior to analysis, the peptides were resuspended in 60 µL MS sample buffer (0.1% formic acid, 3% ACN in water) before subjected to liquid chromatography–mass spectrometry (LC–MS) analysis as previously described (4) .

LC-MS analysis of TMT samples

Samples were loaded and eluted using an UltiMate 3000 RSLCnano system (Thermo Fisher Scientific) with a 220-min gradient separation method. The eluents were then analyzed and quantified with an Orbitrap Fusion mass spectrometer (Thermo Fisher Scientific). MS1 spectra were specified in 375–1,500-m/z scan range with a resolution of 120,000; peptides isolated for MS2 spectra were fragmented by collision-induced dissociation (CID) (30% collision energy); and synchronous precursor selection was used to isolate up to 10 MS2 ions for the MS3-based quantification through high-energy collision-induced dissociation (HCD) (65% collision energy) (6).

Proteomic analysis

Proteomic analysis was performed with the processing software Proteome Discoverer 2.4 (Thermo Fisher Scientific). Peptide sequences were identified by matching theoretical spectra derived from proteome databases with experimental fragmentation patterns via the SEQUEST HT algorithm (7). Fragment tolerances were set to 0.6 Da, and precursor mass tolerances set to 10 ppm with one missed cleavage site allowed. Carbamidomethyl (C, +57.02146) and TMT-tag (K residue and N-terminus of peptide, +229.1629 for 10plex) were specified as static modifications while oxidation (M, +15.994915) was specified as variable. Spectra were searched against the *Homo sapiens* proteome database (Uniprot, 2018, 42,358 sequences) using a false discovery rate of 1% (Percolator) (8, 9). MS3 peptide quantitation was performed with a mass tolerance of 20 ppm. The final list of reported proteins was required to have at least two unique peptides. For enrichment experiments, abundances in each channel were normalized to the endogenous SLC15A4 level. TMT ratios obtained by Proteome Discoverer were transformed with log_2_(x), and p-values were calculated via a Student’s two-tailed t-test with two biological replicates (significant if p < 0.05). Reactome analysis was performed on the list of DMSO versus AJ2-30 proteins (P < 0.05) using Metascape (https://metascape.org/) with the default statistical cutoffs (10).

Immunoblotting and Immunoprecipitation

Immunoblotting and immunoprecipitation were performed as previously described (4). Briefly, treated cells were washed once with ice-cold PBS and lysed in NP-40 buffer (10mM Tris-HCl, pH 7.4; 150mM NaCl, 1% NP-40, 1mM EDTA) supplemented with 1× Halt protease and phosphatase inhibitor for 10 min on ice. Lysates were centrifuged at 13,000 × *g* for 10 min at 4°C, and supernatants were transferred into new tubes. Protein concentrations were determined by the BioRad DC protein assay (5000112). 15 μg proteins were mixed with 4×LDS sample buffer and separated by SDS-PAGE and transferred to a PVDF membrane (Millipore), blocked with SuperBlock blocking buffer (Thermo Fisher, 37536) or 3% BSA in TBST, sequentially incubated with the corresponding primary and secondary antibodies, and visualized with a Bio-Rad ChemiDoc Imaging System.

For immunoprecipitation, cells were resuspended in E1C buffer (50 mM HEPES, pH 7.4; 250mM NaCl, 5 mM EDTA, 0.3% (w/v) CHAPS) with 1x protease inhibitor and sonicated twice (15 msec on, 40 msec off, 15% amplitude, 1 sec total on) on ice. Lysates were cleared by centrifugation. 1 μg antibody or isotype control (Biolegend) were added to each sample and rotated at 4°C overnight followed by incubating with 20 μl protein A/G magnetic beads (MCE) for 1 h. The beads were washed three times with the lysis buffer, proteins eluted with 2× sample buffer at 70°C for 20 min, and resolved by SDS-PAGE. For immunoprecipitation including SLC15A4 blots, beads were eluted by 2× sample buffers at 37°C for 40 min to avoid aggregation.

Nuclear/cytosol fractionation assay

Nuclear-cytoplasm cell fractionation was performed using the REAP method (11, 12). Briefly, cells were harvested, washed once with ice-cold PBS, and lysed in 100 µL cytosolic buffer (PBS containing 0.1% [w/v] NP-40) supplemented with protease inhibitors. Lysates were gently resuspended by pipetting five times with P200 tips and subjected to a brief centrifugation (“pop-spin”; 10,000 × *g* for 10 s). The supernatant, containing the cytosolic fraction, was transferred to a new tube. The pellet, containing nuclei, was lysed in 50 µL RIPA buffer supplemented with protease inhibitors, incubated on ice for 5 min, and centrifuged at 10,000 rpm for 5 min at 4°C. The resulting supernatant was collected as the nuclear fraction. Fractionation purity was assessed by immunoblotting for GAPDH (cytosolic marker) and Lamin B (nuclear marker).

Quantitation of cytokines levels by Enzyme-linked immunosorbent assay (ELISA)

Cytokine levels were quantified by ELISA using Human TNF (BioLegend), IL-6 (BioLegend), and IFN-β (InvivoGen) kits according to the manufacturers’ instructions. Undiluted culture supernatants were used for all measurements. Absorbance or luminescence was measured using a CLARIOstar plate reader (BMG Labtech). All experiments were performed with at least three independent biological replicates.

RNA Isolation and Quantitative Real-Time PCR

Cells were collected, washed once with ice-cold PBS, and total RNA was isolated using the GeneJET RNA Isolation Kit (Thermo Fisher Scientific) according to the manufacturer’s instructions. Reverse transcription was performed using the PrimeScript™ RT Reagent Kit (Perfect Real Time; Takara) with a combination of random hexamer and oligo(dT) primers. Quantitative real-time PCR was carried out using PowerUp™ SYBR™ Green Master Mix (Thermo Fisher Scientific). Gene-specific primer sequences are listed in *Table S3*. Relative gene expression was calculated using the ΔΔCt method, with normalization to GAPDH and the unstimulated control group. Each sample was analyzed in technical duplicates, and all experiments were performed with three independent biological replicates.

Immunofluorescence

Immunofluorescence staining was performed using the BD Cytofix/Cytoperm™ Fixation/Permeabilization Kit (554714) with minor modifications. Briefly, cells were seeded onto coverslips coated with 0.02% poly-L-lysine (Sigma, P1274) at room temperature for 15 min, fixed with BD fixation/permeabilization solution for 10 min at room temperature, and permeabilized with 1× BD Perm/Wash™ (PW) buffer for 30 min at room temperature. Primary antibodies were diluted at the indicated ratios in PW buffer and incubated with cells overnight at 4°C. After three washes with PW buffer, cells were incubated with Alexa Fluor–conjugated secondary antibodies (1:500) for 1 h at room temperature. Nuclei were counterstained with NucBlue™ Live Cell Stain (Thermo Fisher Scientific, R37606) for 5 min, and coverslips were mounted using ProLong™ Diamond Antifade Mountant (Thermo Fisher Scientific, P36965).

Confocal images were acquired on a Zeiss LSM 710 microscope using a 100× oil-immersion objective (NA = 1.40), with the pinhole set to 1 Airy unit for each channel. Images were processed using ImageJ software. For colocalization analysis, 25–50 cells were randomly selected per condition, and channel-specific thresholds were applied. Colocalization was quantified using ZEN 2011 SP7 software (version 14.0.5.201) and expressed as a colocalization coefficient calculated as the number of colocalized pixels between proteins A and B divided by the total number of pixels for protein A.

Flow cytometry

For SLC15A4 and TLR9 expression analysis, intracellular staining was performed using the BD Cytofix/Cytoperm™ Fixation/Permeabilization Kit with minor modifications. Briefly, 10^5^ cells in 96-well plates were fixed with BD fixation/permeabilization solution for 10 min at room temperature and permeabilized with 1× PW buffer for 30 min at room temperature. Primary antibodies (anti-SLC15A4, 1:1500; anti-TLR9, 1:200) were diluted in PW buffer and incubated with cells for an hour at room temperature. After one wash with PW buffer, cells were incubated with Alexa Fluor–conjugated secondary antibodies (1:500) for 30 min at room temperature. Cells were then washed with PW buffer twice, and resuspended in FACS buffer (PBS, 2% FBS, 1mM EDTA) prior to analysis. For detection of S6 phosphorylation, cells were fixed in 4% paraformaldehyde for 10 min, permeabilized with 1× True-Phos™ Perm Buffer (BioLegend, 425401) for 1h at −20 °C, and washed twice with FACS buffer. Cells were then incubated with Alexa Fluor–conjugated antibodies for 1 h at room temperature, washed, resuspended in FACS buffer, and analyzed by flow cytometry. Flow cytometry experiments were performed in the Scripps Research Flow Core using a NovoCyte flow cytometer (ACEA Biosciences, Inc.). FACS data was analyzed using the software FlowJo v10.8.0 (BD Biosciences).

Predicting the interaction structure between SLC15A4 and the LAMTOR regulatory complex.

Canonical sequences of SLC15A4, LAMTOR1, LAMTOR2, LAMTOR3, LAMTOR4, and LAMTOR5 (UniProt accession numbers Q8N697, Q6IAA8, Q9Y2Q5, Q9UHA4, Q0VGL1, and O43504, respectively) were used for structural predictions. Structural predictions were performed using AlphaFold 3 (13). Repeats were run with different seeds each generating five independent models. A pose of interest where LAMTOR1’s alpha helix protruded into the SLC15A4 binding site was recapitulated in seed 3 of the AlphaFold 3 diffusion sampling, with 3 out of the five poses exhibiting this complex geometry (**Fig. S9**). Despite having a prediction confidence near the model’s confidence boundary, this candidate was selected for subsequent analyses.

Statistical analysis

Western blot signals were quantified using ImageJ from a minimum of two biological replicates. Phosphorylated protein intensities were first normalized to the corresponding total (unphosphorylated) protein and then normalized to untreated (0 h) samples within each experiment. For nuclear translocation assays, nuclear protein levels were first normalized to Lamin B and subsequently normalized to untreated (0 h) samples. Cytokine production was measured by ELISA using three biological replicates. RT–PCR analyses were performed in two technical replicates, with all experiments repeated in three independent biological replicates. Statistical significance at each time point was determined using a two-tailed Student’s t-test or One-way ANOVA followed by Sidak’s multiple comparisons test. For IP–MS experiments, statistical analysis was performed using a two-tailed Student’s t-test with two biological replicates, with significance defined as p < 0.05. Colocalization coefficients are presented as box plots, where the center line indicates the median, the box represents the 25th–75th percentiles, and whiskers denote minimum and maximum values. Statistical comparisons were performed using a two-tailed Student’s t-test. *, p ≤ 0.05; **, p ≤ 0.01; ***, p ≤ 0.001; ****, p ≤ 0.0001; ns, not significant.

Resource availability

Lead contact and materials availability

Schematic figures

Figures have been created in BioRender (Created in <https://BioRender.com>)

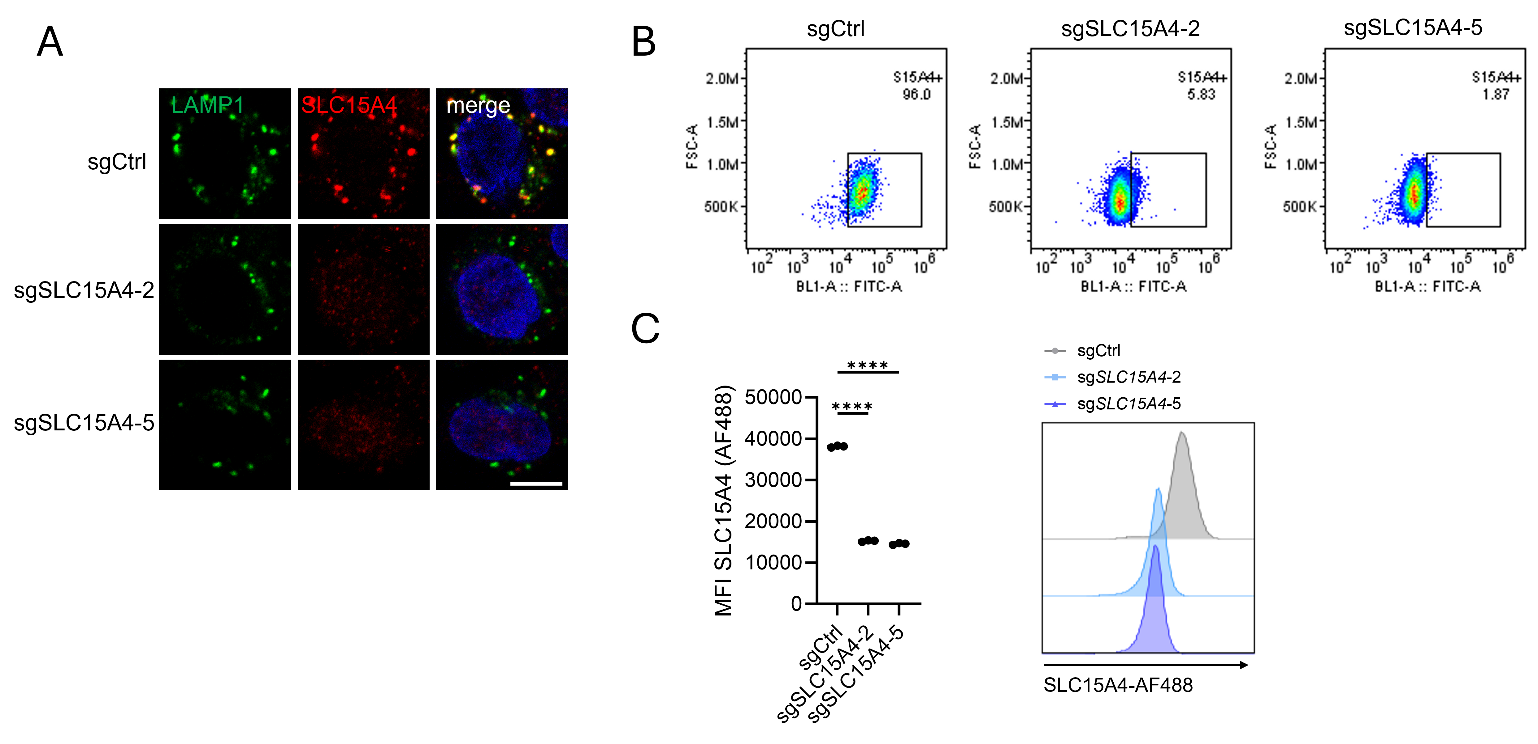

Fig. S1. Validation of SLC15A4-knockout CAL-1 cells

(A) Immunostaining of endogenous SLC15A4 in CAL-1 cells. Cells were co-stained with LAMP1 as a lysosomal marker. sgCtrl, single guide RNA targeting Renilla luciferase; sgSLC15A4-2 and sgSLC15A4-5, two independent SLC15A4-knockout clones. Scale bar, 5 μm.

(B-C) SLC15A4 expression level in control and SLC15A4-knockout cells were assessed by flow cytometry and calculated as the mean fluorescence intensity (MFI). Statistical comparisons were performed using One-way ANOVA test. ****, p ≤ 0.0001.

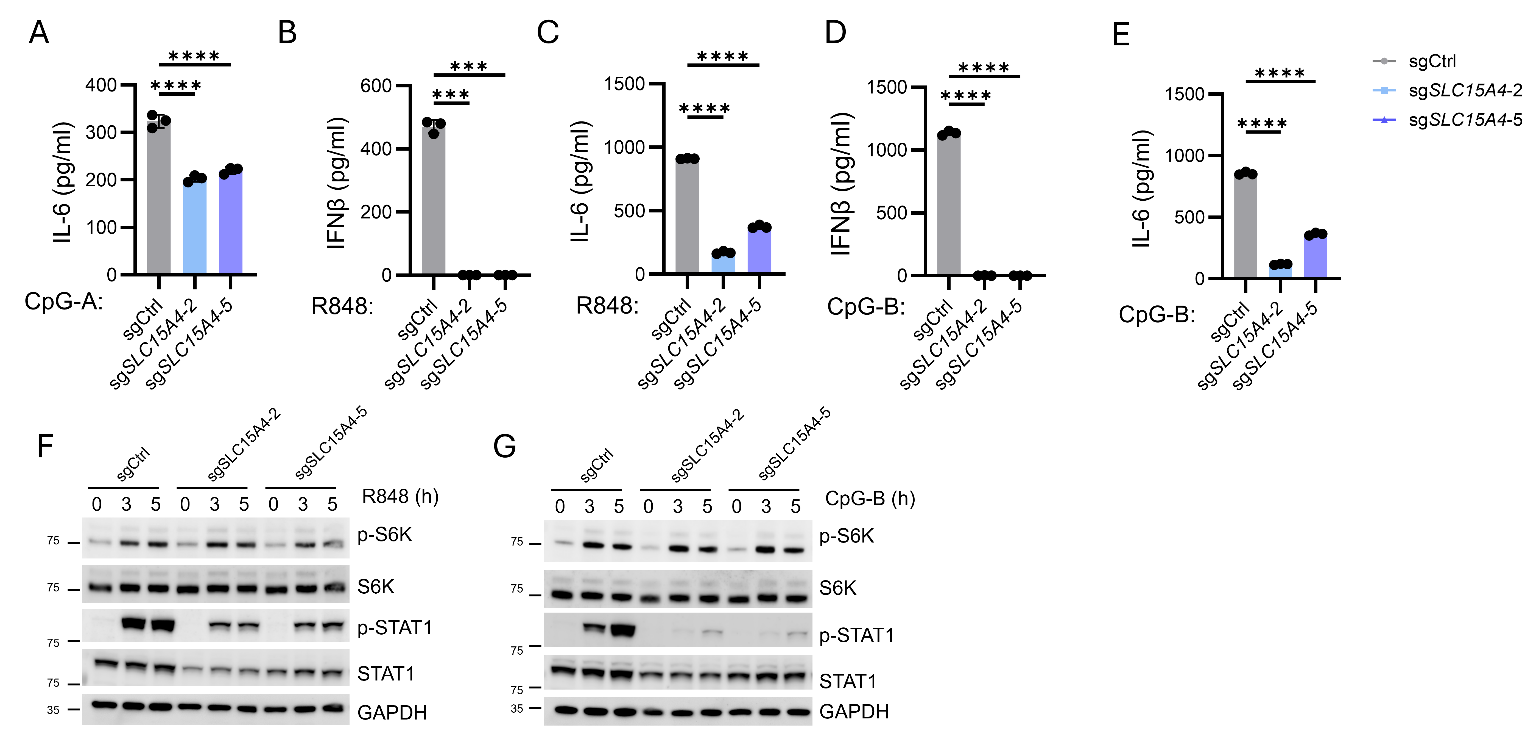

Fig. S2. SLC15A4 mediates endolysosomal TLR-induced mTOR and cytokine production via multiple circuits, related to Fig. 1.

(A-E) Cytokine production in gene-edited CAL-1 cells with either control or SLC15A4 sgRNAs following stimulation with (A) CpG-A, (B-C) R848, or (D-E) CpG-B for 24 h. Data are shown as mean ± s.d. from n = 3 biological replicates.

(F, G) Immunoblots from gene-edited CAL-1 cells with either control or SLC15A4 sgRNAs stimulated with R848 or CpG-B for 0–5 h. p-, phosphorylated.

Statistical comparisons were performed using a one-way ANOVA test. ***, p ≤ 0.001; ****, p ≤ 0.0001.

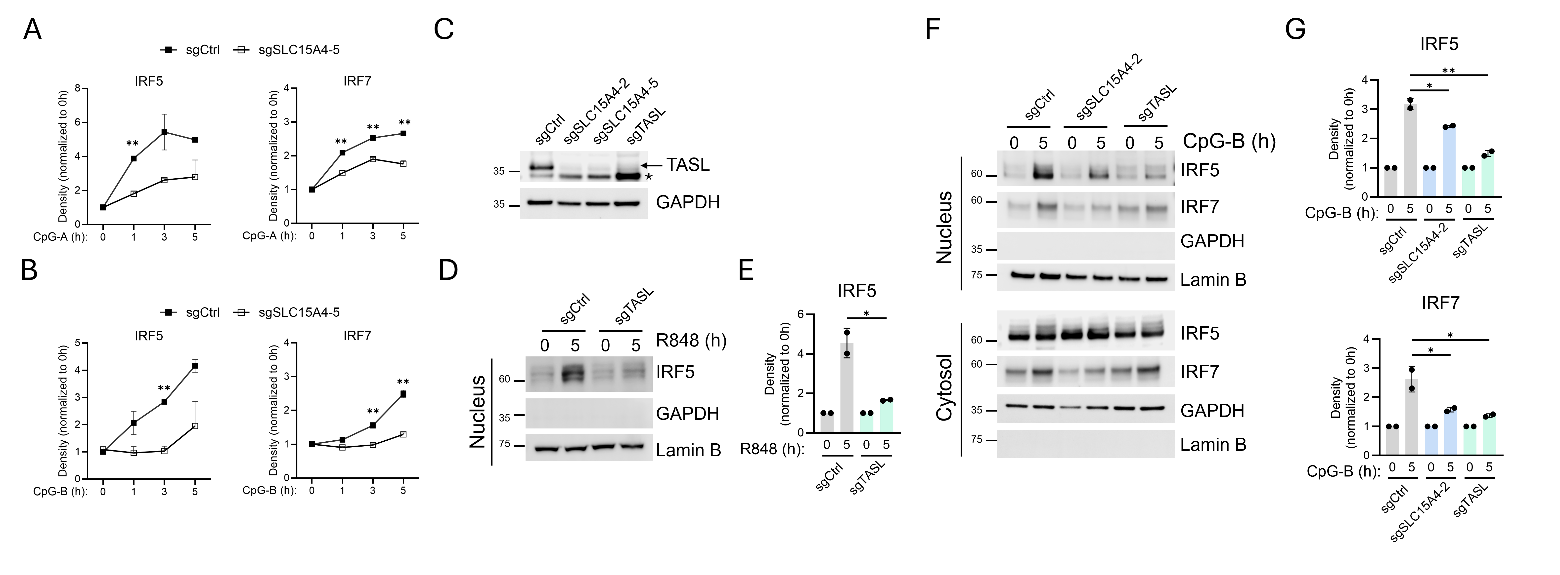

Fig. S3. SLC15A4 mediates TLR7-9-induced IRF5/7 activation, related to Fig. 1.

(A, B) Quantification of band intensities from Fig. 1F and 1G.

(C) Immunoblots of SLC15A4 and TASL-knockout CAL-1 cells with TASL antibody. * indicates non-specific band.

(D, E) Nuclear fractionation from control and TASL-knockout CAL-1 cells stimulated with R848. Quantification of nuclear IRF5 was shown.

(F, G) Nuclear fractionation and quantification from control, SLC15A4-knockout (clone 2) and TASL-knockout CAL-1 cells stimulated with CpG-B. GAPDH, cytosolic fraction control. Lamin B, nuclear fraction control.

Statistical significance at each time point was determined using a two-tailed Student’s t-test or a One-way ANOVA test (for G). *, p ≤ 0.05; **, p ≤ 0.01; ns, not significant.

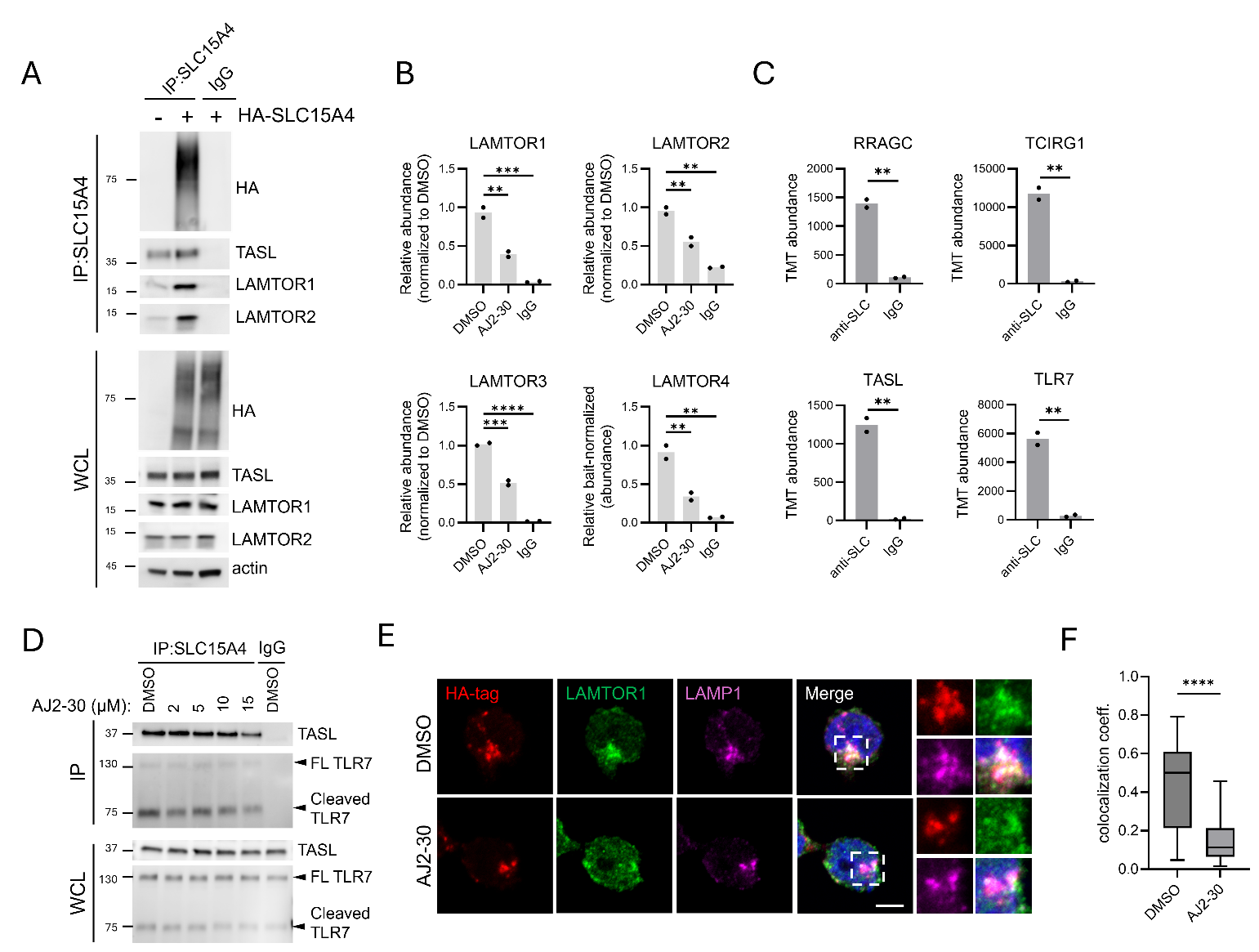

Fig. S4. SLC15A4 directly engages the Ragulator-Rag complex, related to Fig. 2.

(A) Immunoprecipitation of endogenous SLC15A4 in HA-tagged SLC15A4 expressed CAL-1 cells and analyzed by immunoblot. WCLs, whole cell lysates. IP, immunoprecipitation.

(B) Relative enriched-TMT abundance (normalized to SLC15A4 bait levels) of LAMTOR1-4 from Fig. 2B-C.

(C) TMT abundance (normalized to bait level) of other SLC15A4-enriched proteins from Fig. 2B-C.

(D) Immunoprecipitation of endogenous SLC15A4 in CAL-1 cells followed by immunoblot analysis. WCLs, whole cell lysates; IP, immunoprecipitation. FL, full length. Data are representative of two independent experiments.

(E) Immunostaining of overexpressed SLC15A4 and endogenous LAMTOR1 in CAL-1 cells. Cells were treated with AJ2-30 or DMSO for 1 h and co-stained with LAMP1 as a lysosomal marker. Zoomed in regions are indicated by white dashed boxes. Scale bar, 5 μm.

(F) Quantification of the colocalization coefficient of HA-SLC15A4 with LAMTOR1. Images are representative of two independent experiments.

For S4B-C, statistical analysis was performed using two-tailed Student’s test. For S4F, Colocalization coefficients are presented as box plots, where the center line indicates the median, the box represents the 25th–75th percentiles, and whiskers denote minimum and maximum values. Statistical comparisons were performed using a two-tailed Student’s t-test. **, p ≤ 0.01; ***, p ≤ 0.001; ****, p ≤ 0.0001.

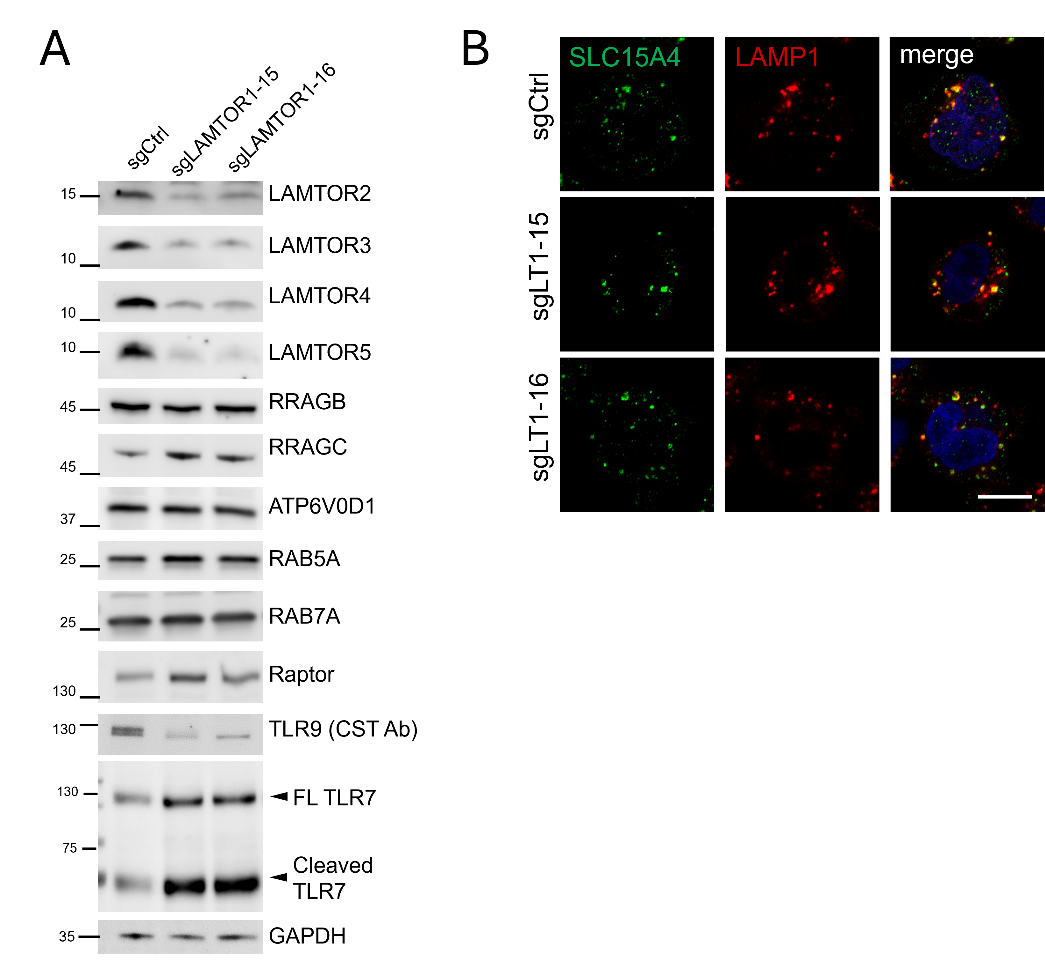

Fig. S5. Characterization of LAMTOR1-KO cells, related to Fig. 3.

(A) Immunoblots of indicated proteins in control and LAMTOR1-knockout CAL-1 cells. Note that the anti-TLR9 antibody (Cell Signaling Technology) only recognizes full-length TLR9 but not the cleaved TLR9 by immunoblot.

(B) Immunostaining of endogenous SLC15A4 in gene-edited CAL-1 cells with either LAMTOR1 or control sgRNAs. Cells were co-stained with LAMP1 as a lysosomal marker. Scale bar, 5 μm. LT1, LAMTOR1.

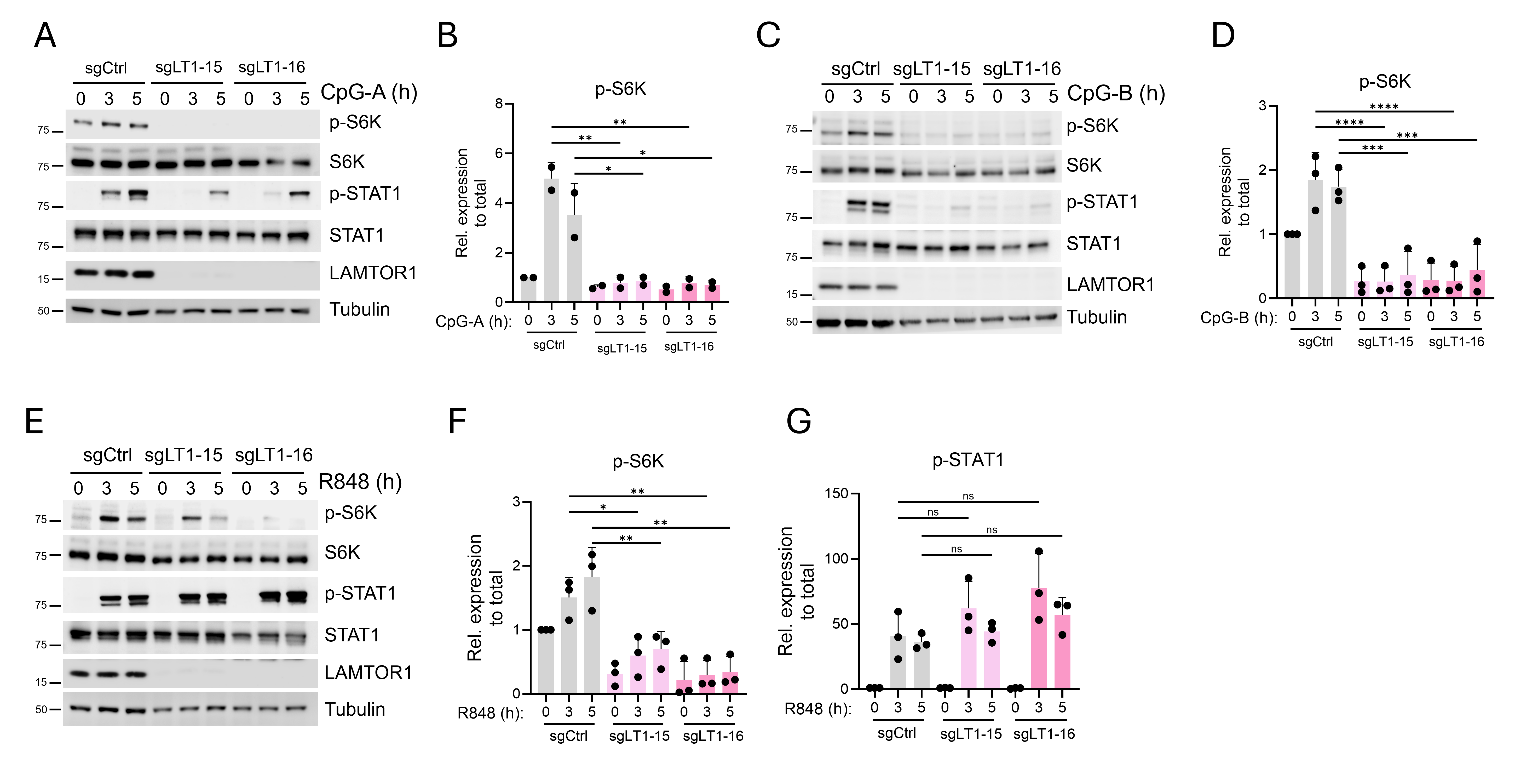

Fig. S6. *LAMTOR1* loss differentially impacts endolysosomal TLR mediated signaling, related to Fig. 3.

(A-G) Immunoblot analysis and quantification in gene-edited CAL-1 cells with either LAMTOR1 or control sgRNAs following (A, B) CpG-A, (C, D) CpG-B, (E-G) R848 stimulation for 0–5 h. p-, phosphorylated. LT1, abbreviation for LAMTOR1.

Statistical significance at each time point was determined using a One-way ANOVA test. *, p ≤ 0.05; **, p ≤ 0.01; ***, p ≤ 0.001; ****, p ≤ 0.0001; ns, not significant.

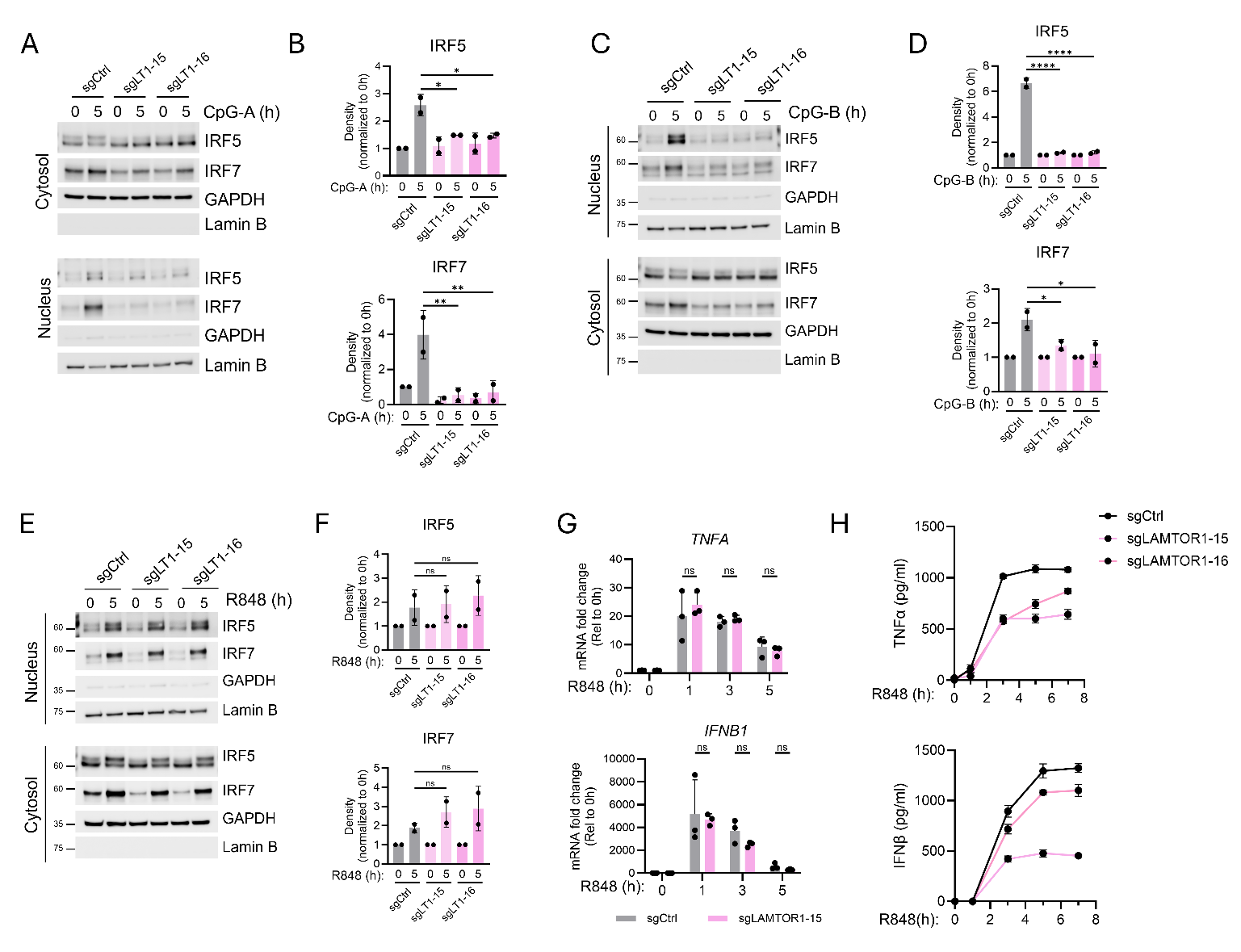

Fig. S7. Characterization of LAMTOR1-KO cells upon TLR7-9 stimulation, related to Fig. 3.

(A-F) Nuclear fractionation and quantification of gene-edited CAL-1 cells with either LAMTOR1 or control sgRNAs stimulated with (A, B) CpG-A (C, D) CpG-B, and (E, F) R848. GAPDH, cytosolic fraction control. Lamin B, nuclear fraction control.

(G) *TNF* and *IFNB1* mRNA levels in gene-edited CAL-1 cells with either LAMTOR1 or control sgRNAs following R848 stimulation for 0-5 h, measured by RT-qPCR. Fold changes were calculated relative to the unstimulated group. Data are presented as mean ± s.d. from n = 3 biological replicates.

(H) Cytokine production in gene-edited CAL-1 cells with either LAMTOR1 or control sgRNAs following stimulation with R848 for 0-7h. Data are presented as mean ± s.d. from n = 3 biological replicates.

Statistical significance at each time point was determined using a two-tailed Student’s t-test (for J) or a One-way ANOVA test. *, p ≤ 0.05; **, p ≤ 0.01; ****, p ≤ 0.0001; ns, not significant.

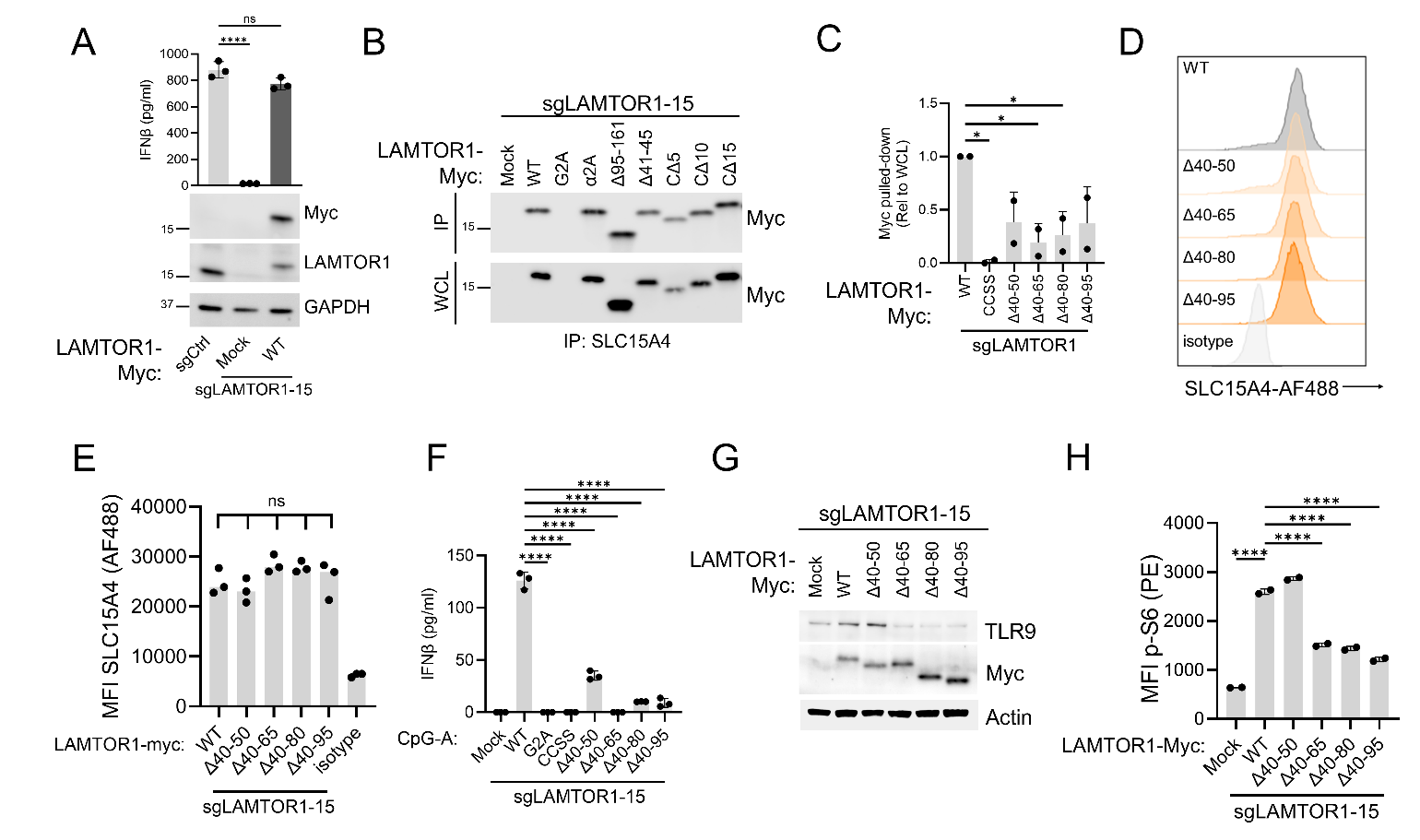

Fig. S8. Interaction of truncated LAMTOR1 and SLC15A4, related to Fig. 4.

(A) Expression of LAMTOR1 and IFNβ production in control, LAMTOR1-knockout, and wild-type LAMTOR1 reconstituted CAL-1 cells following stimulation with CpG-B for 24 h. Data are shown as mean ± s.d. from n = 3 biological replicates.

(B) Immunoprecipitation of endogenous SLC15A4 with indicated LAMTOR1 mutants was performed in LAMTOR1-knockout CAL-1 cells reconstituted with LAMTOR1 mutants and analyzed by immunoblot. α2A, Y88/L92/V94/L95/L99/W102A; CΔ5, C∆10, or C∆15 indicates deletion of C-terminal 5, 10, or 15 amino acids of LAMTOR1, respectively, as previously described (14). WCLs, whole cell lysates. IP, immunoprecipitation.

(C) Quantification of immunoprecipitation results in Fig 4C.

(D, E) SLC15A4 levels in LAMTOR1-knockout CAL-1 cells reconstituted with LAMTOR1 mutants were assessed by flow cytometry and depicted as the mean fluorescence intensity (MFI).

(F) IFN-β production in LAMTOR1-knockout CAL-1 cells reconstituted with LAMTOR1 mutants following stimulation with CpG-A for 24 h.

(G) Immunoblots of anti-TLR9 (Cell Signaling Technology) and anti-Myc-tag in LAMTOR1-knockout CAL-1 cells reconstituted with LAMTOR1 mutants.

(H) Phosphorylated levels of S6 upon CpG-B stimulation in in LAMTOR1-knockout CAL-1 cells reconstituted with LAMTOR1 mutants were assessed by flow cytometry and depicted as the mean fluorescence intensity (MFI).

Results were compared with LAMTOR1-WT cells. Statistical significance was determined using a One-way ANOVA test. *, p ≤ 0.05; **, p ≤ 0.01; ns, not significant.

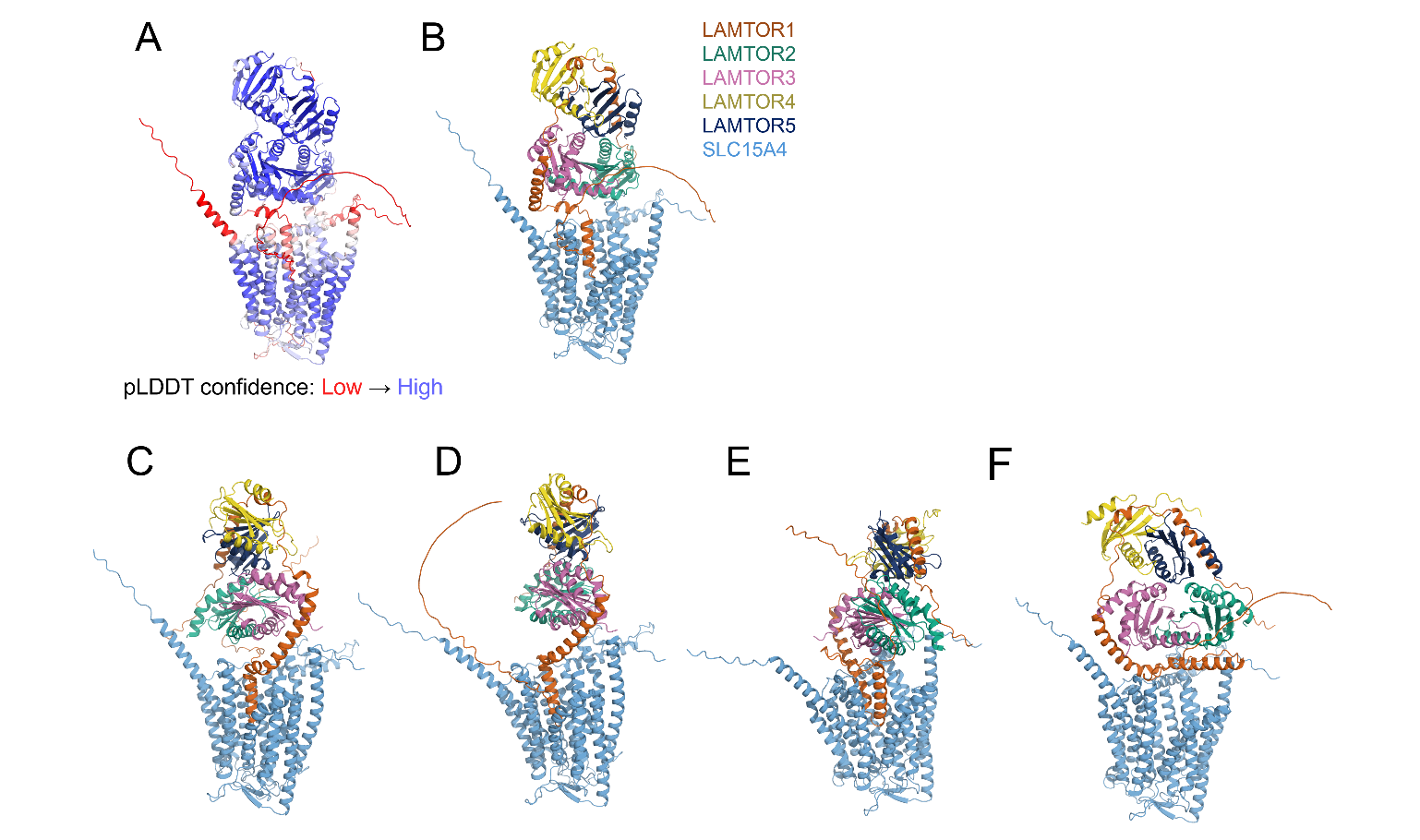

Fig. S9. Structural predictions of the SLC15A4 hexametric complex, related to Fig. 4A.

(A) Representative AlphaFold3 predicted structure colored by pLDDT (predicted Local Distance Difference Test) confidence (red = low, blue = high).

(B-F) Cartoon representations of the top-ranked models from five independent AlphaFold3 diffusion samples (seed 3, samples 0-4; panels A–E). The complex comprises SLC15A4 (light blue, transmembrane), LAMTOR1 (orange), LAMTOR2 (green), LAMTOR3 (magenta), LAMTOR4 (yellow), and LAMTOR5 (dark blue). Structures were visualized in PyMOL and colored by chain.

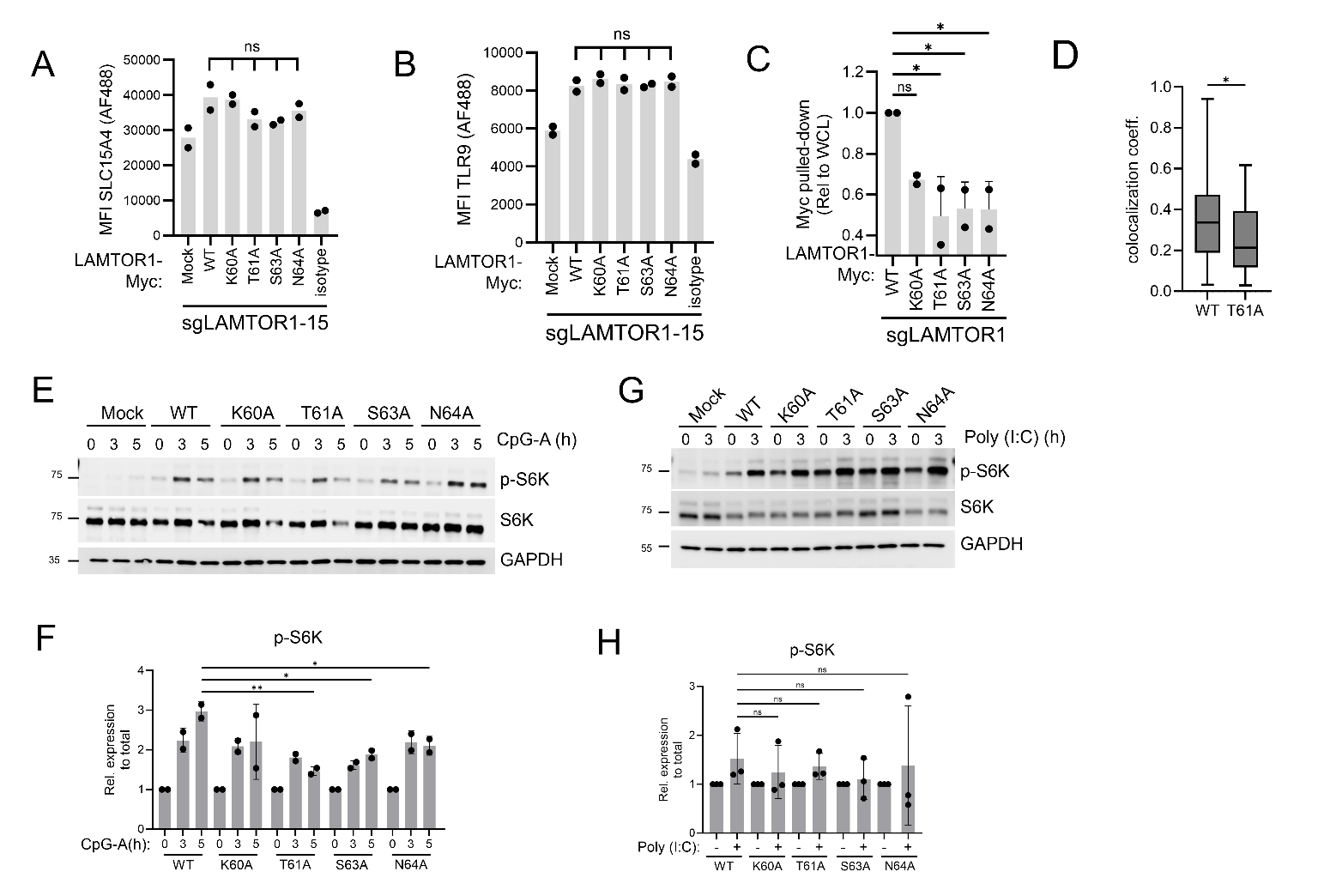

Fig. S10. Characterization of LAMTOR1 mutants, related to Fig. 4.

(A-B) SLC15A4 (A) or TLR9 (B) protein levels in LAMTOR1-knockout CAL-1 cells reconstituted with the indicated WT and mutant Myc-LAMTOR1 constructs. Samples were assessed by flow cytometry and depicted as the mean fluorescence intensity (MFI).

(C) Quantification of immunoprecipitation results in Fig 4D.

(D) Quantification of SLC15A4 colocalization with WT and mutant Myc-LAMTOR1 constructs in Fig 4E.

(E, F) Immunoblots and quantification of LAMTOR1-knockout CAL-1 cells reconstituted with WT and mutant Myc-LAMTOR1 constructs upon stimulation with CpG-A for 0–5h. p-, phosphorylated.

(G, H) Immunoblots and quantification of LAMTOR1-knockout CAL-1 cells reconstituted with WT and mutant Myc-LAMTOR1 constructs upon stimulation with poly(I:C) for 3h. p-, phosphorylated.

Results were compared with WT Myc-LAMTOR1 cells. Statistical significance was determined using a two-tailed Student’s t-test or One-way ANOVA test (for D). *, p ≤ 0.05; **, p ≤ 0.01; ***, p ≤ 0.001; ****, p ≤ 0.0001; ns, not significant.

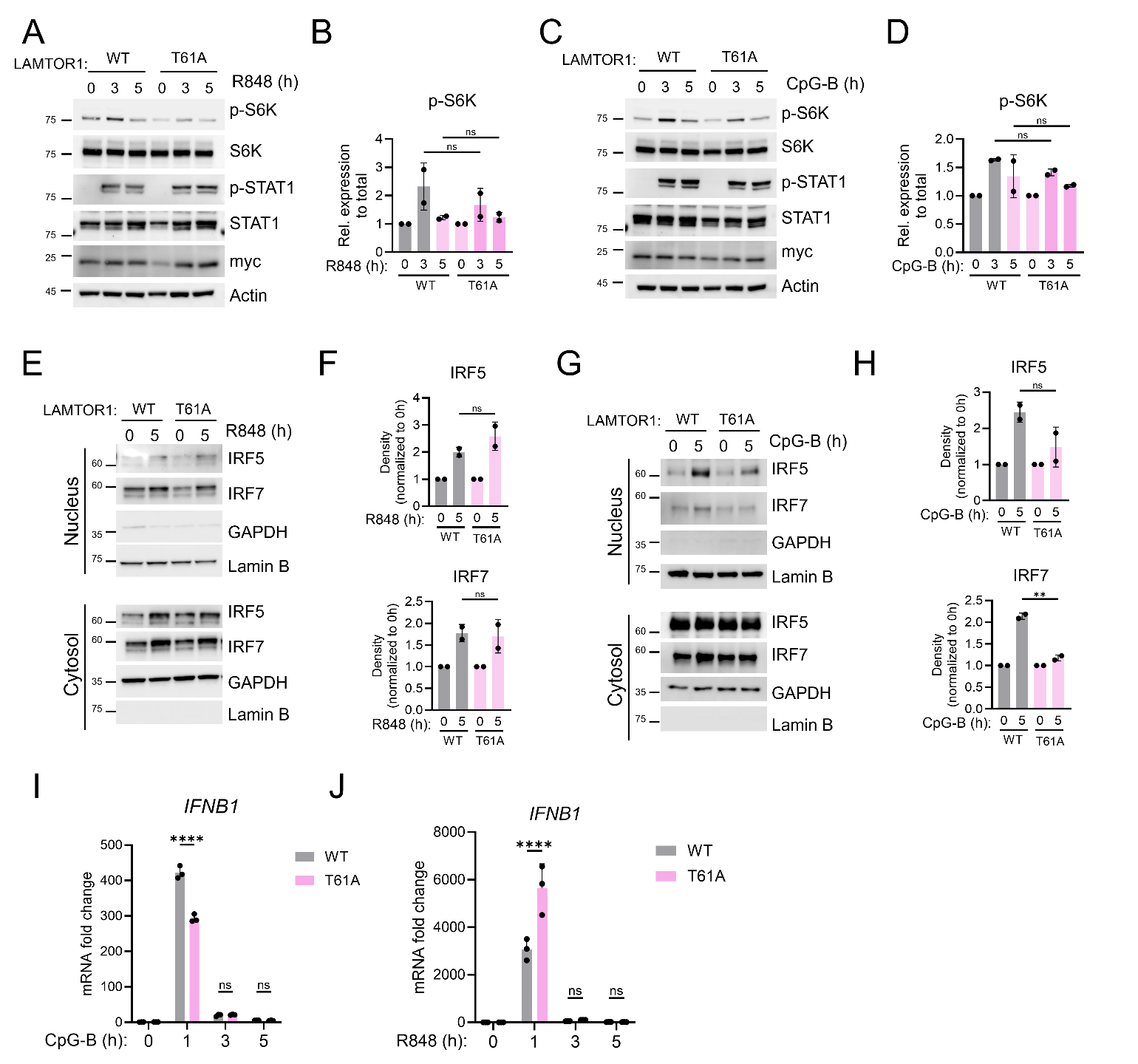

Fig. S11. Characterization of LAMTOR1 T61A mutation in response to TLR agonists, related to Fig. 4.

(A-D) Immunoblots and quantification of LAMTOR1-knockout CAL-1 cells reconstituted with WT and T61A-mutant Myc-LAMTOR1 constructs stimulated with (A, B) R848 or (C, D) CpG-B for 0–5h. p-, phosphorylated.

(E-H) Nuclear fractionation and quantification of LAMTOR1-knockout CAL-1 cells reconstituted with WT and T61A-mutant Myc-LAMTOR1 constructs stimulated with (E, F) R848 or (G, H) CpG-B for 5h. GAPDH, cytosolic fraction control. Lamin B, nuclear fraction control.

(I, J) *IFNB1* mRNA levels of LAMTOR1-knockout CAL-1 cells reconstituted with WT and T61A-mutant Myc-LAMTOR1 constructs stimulated with (I) CpG-B or (J) R848 for 0–5h, measured by RT-qPCR. Fold changes were calculated relative to the unstimulated group. Data are presented as mean ± s.d. from n = 3 biological replicates.

Results were compared with WT Myc-LAMTOR1 cells. Statistical significance was determined using a one-way ANOVA test (for B, D) or a two-tailed Student’s t-test (for F, H). *, p ≤ 0.05; **, p ≤ 0.01; ***, p ≤ 0.001; ****, p ≤ 0.0001; ns, not significant.

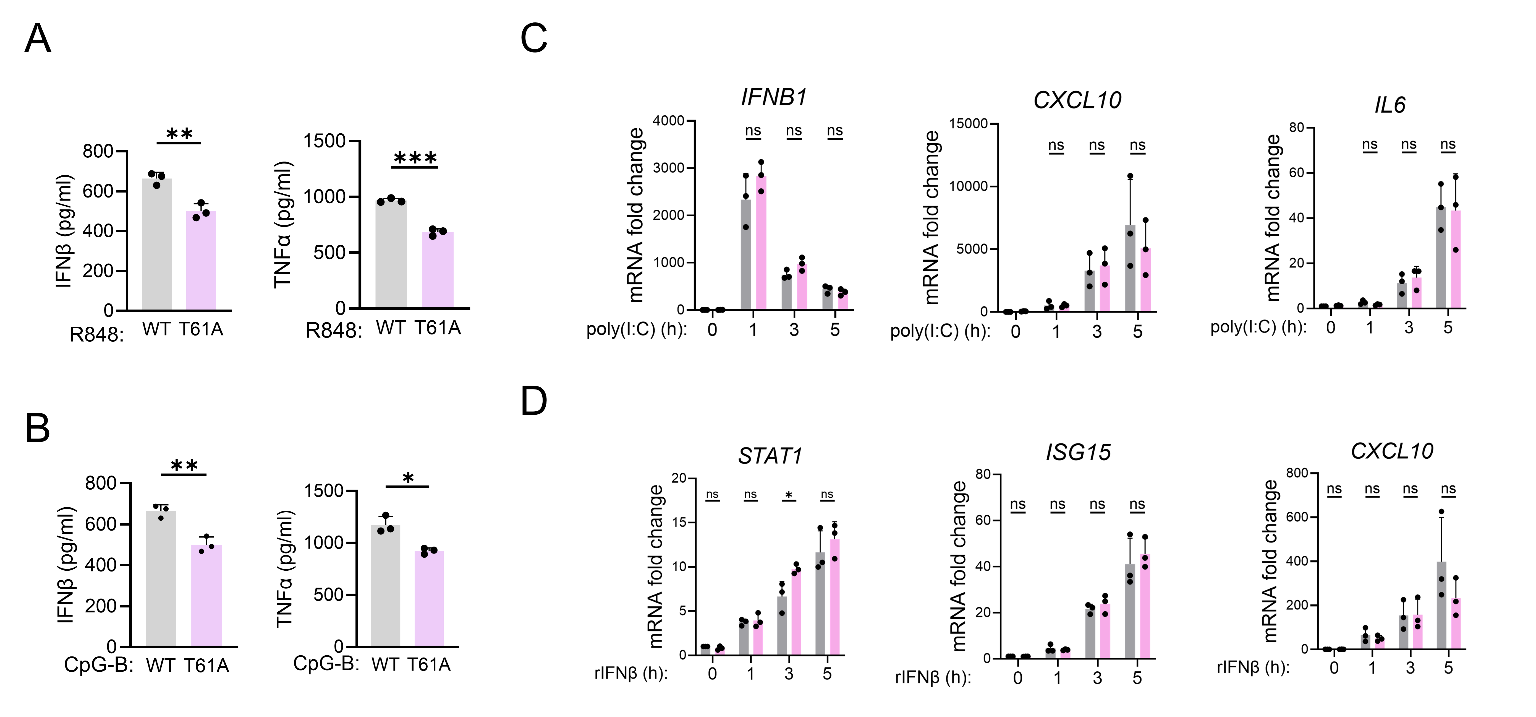

Fig. S12. Profiling of LAMTOR1 T61A mutation in response to TLR agonists, related to Fig. 4.

(A, B) Cytokine production in the indicated CAL-1 cells stimulated with R848 or CpG-B for 24 h.

(C) *IFNB1*, *CXCL10, and IL-6* mRNA levels in the indicated CAL-1 cells following poly(I:C) stimulation, measured by RT–PCR. Fold changes were calculated relative to the unstimulated group.

(D) IFN-stimulated genes STAT1, ISG15, and CXCL10 mRNA levels in the indicated CAL-1 cells following recombinant IFNβ stimulation, measured by RT–PCR. Fold changes were calculated relative to the unstimulated group.

Results were compared with WT Myc-LAMTOR1 cells. Statistical significance was determined using a two-tailed Student’s t-test. *, p ≤ 0.05; **, p ≤ 0.01; ***, p ≤ 0.001; ****, p ≤ 0.0001; ns, not significant.

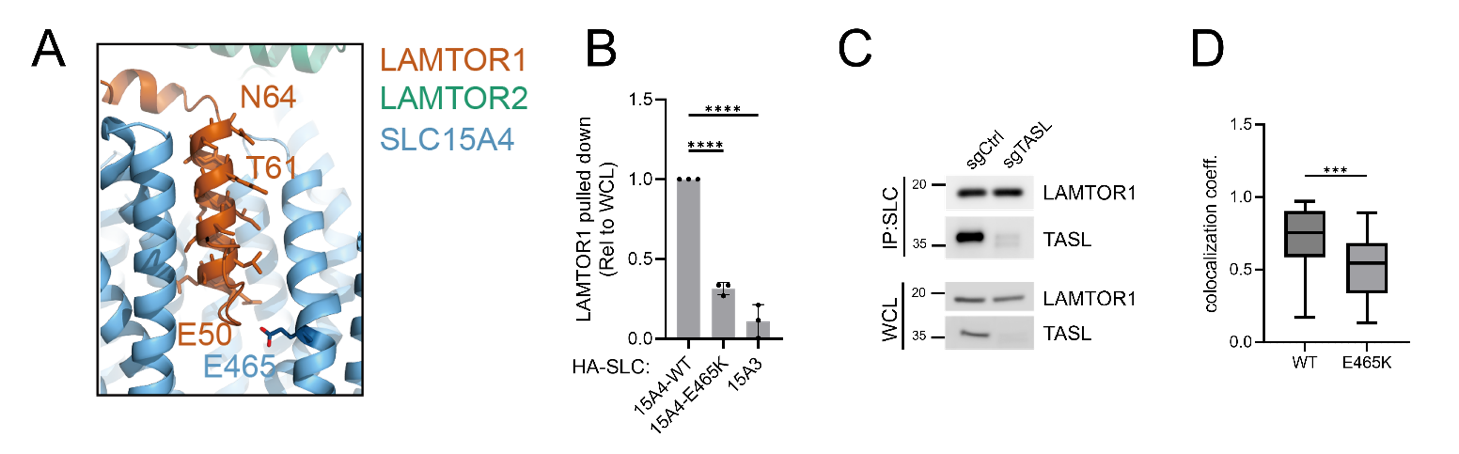

Fig. S13. The SLC15A4 residue E465 is essential for LAMTOR1 association with SLC15A4, related to Fig. 5.

(A) AlphaFold prediction of interactions between SLC15A4 and LAMTOR1. LAMTOR1 α1-helix residues E50 and N64 are highlighted in brown. SLC15A4 residue E465 is colored in blue.

(B) Quantification of immunoprecipitation results in Fig. 5A.

(C) Immunoprecipitation of endogenous SLC15A4 in control and TASL-knockout CAL-1 cells, followed by immunoblot analysis.

(D) Quantification of LAMTOR1 colocalization with WT and E465K-mutant HA-SLC15A4 constructs in Fig. 5B.

Results were compared with WT HA-SLC15A4 cells. Statistical significance was determined using a one-way ANOVA test (For B) or a two-tailed Student’s t-test (For D). ***, p ≤ 0.001; ****, p ≤ 0.0001.

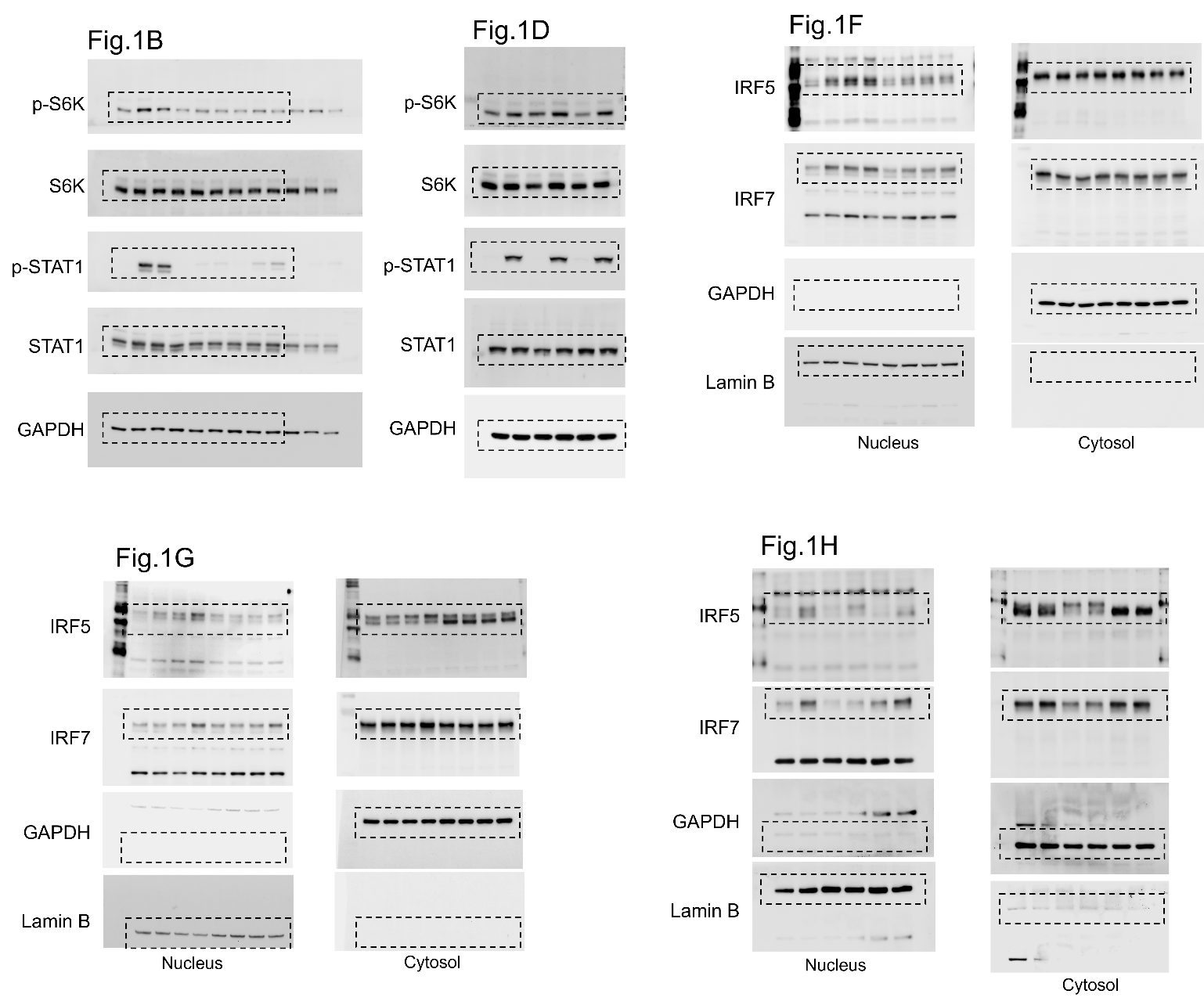

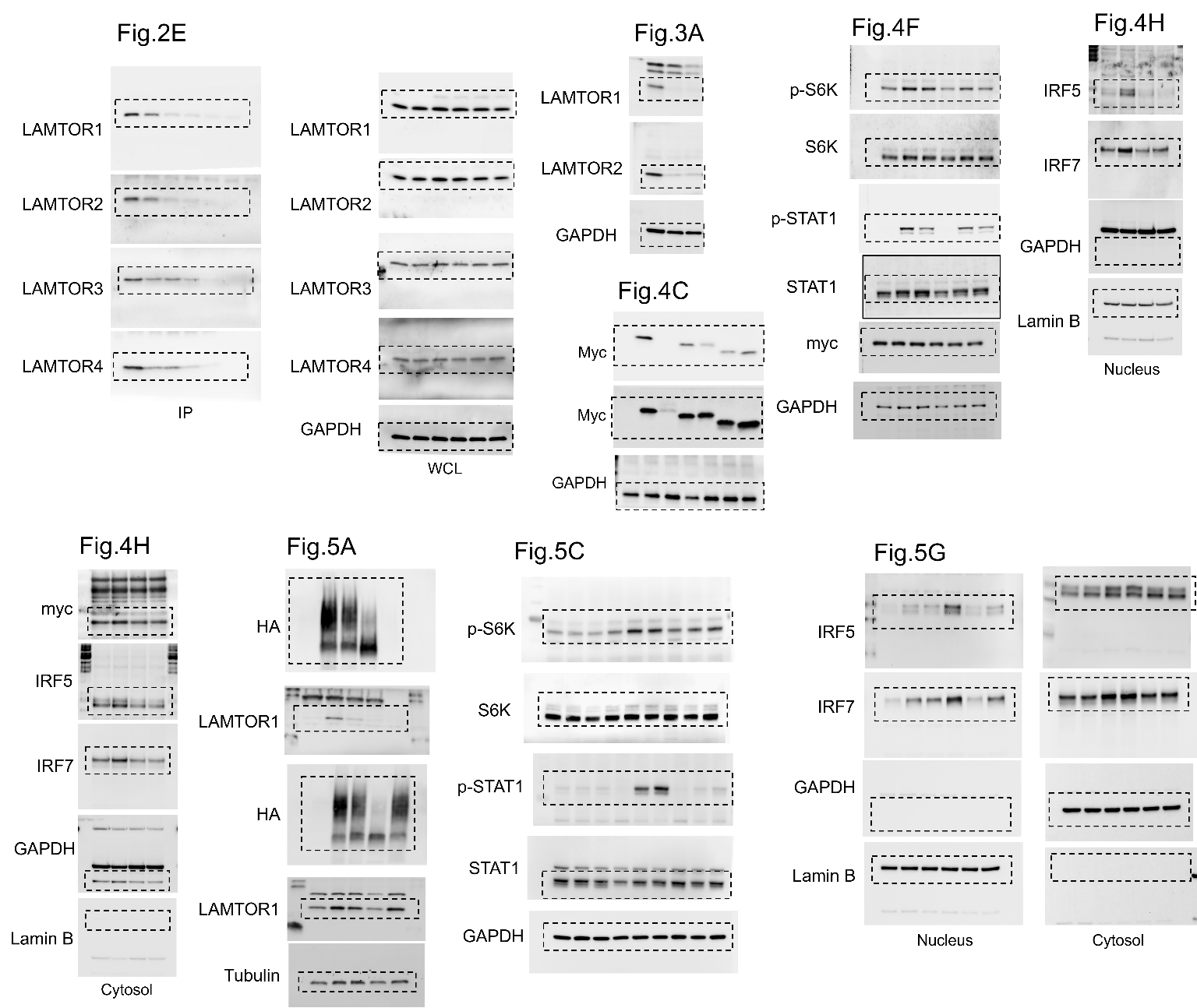

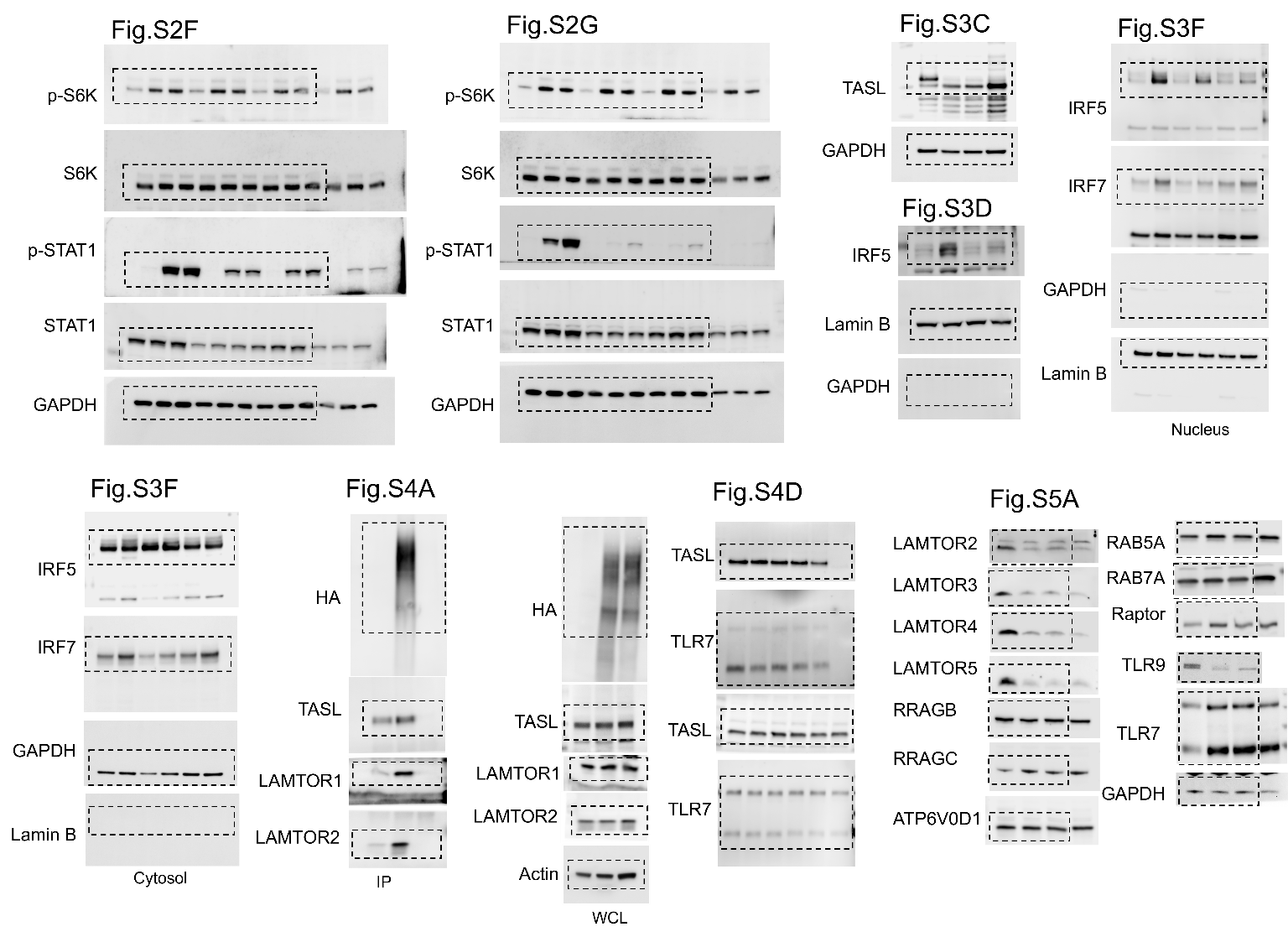

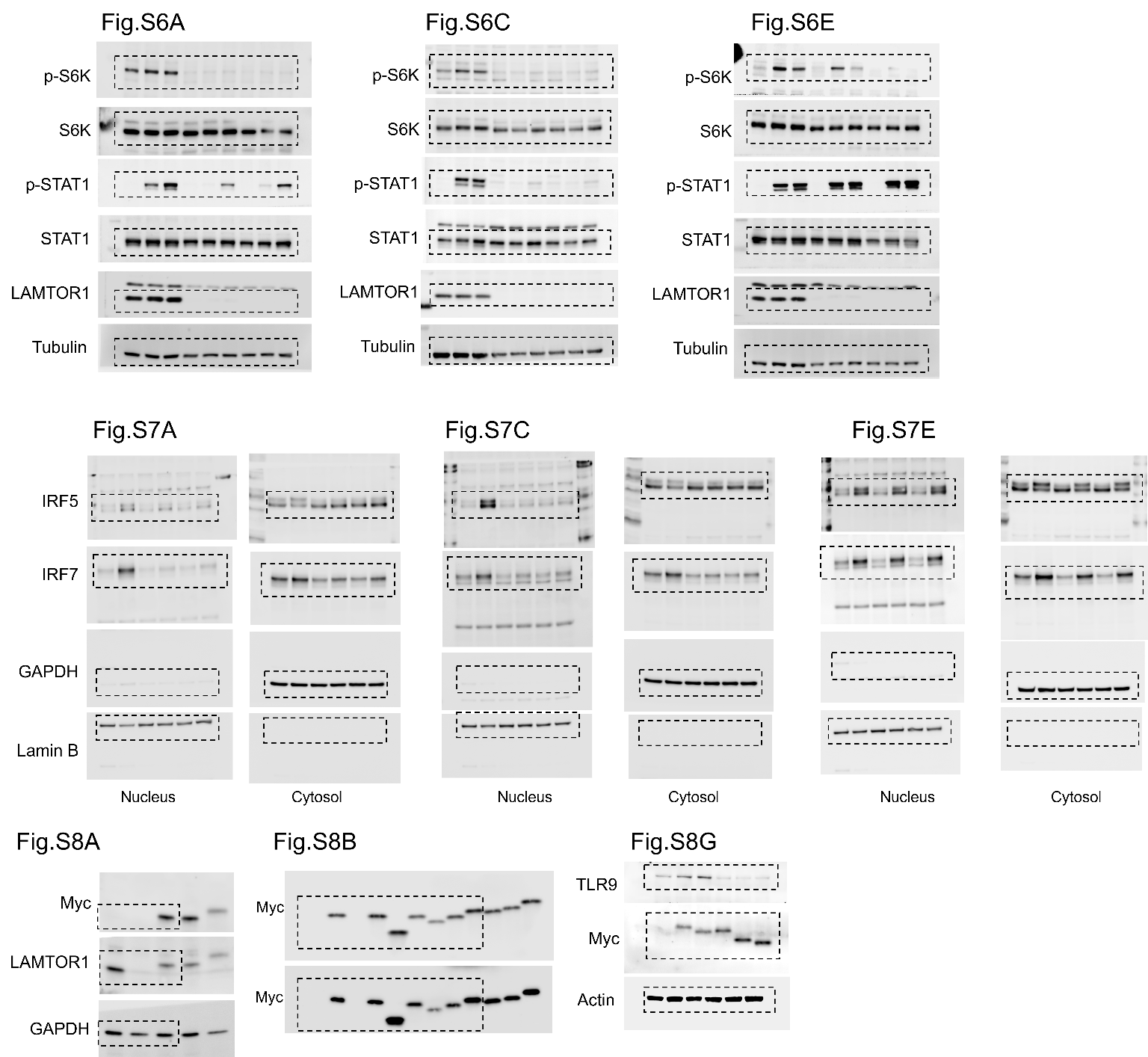

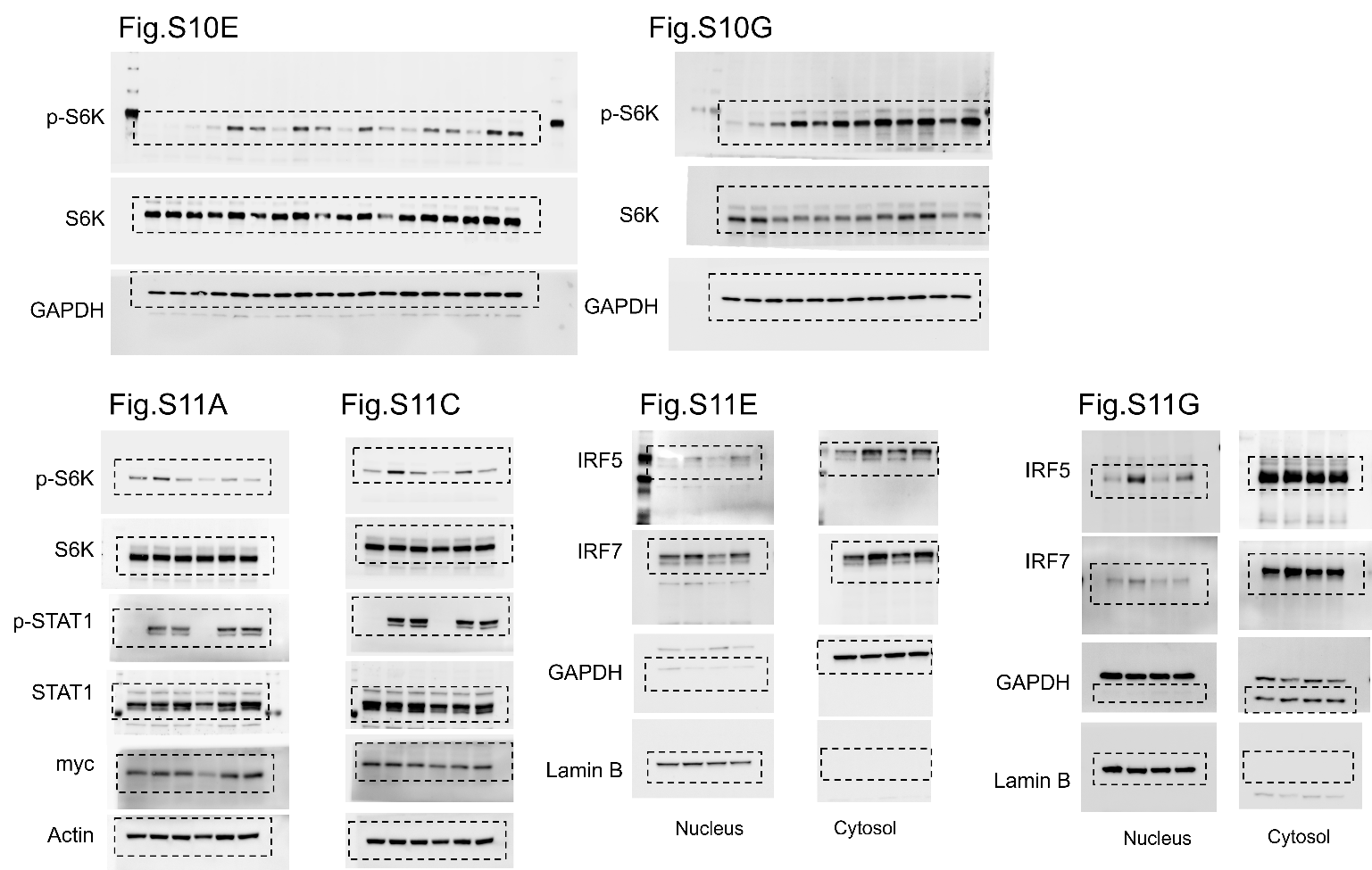

Fig. S14. Uncropped western blots of main figures and supplementary information.

**Table S1. List of sgRNA for CRISPR/Cas9 gene-editing (5′ to 3′ orientation)**

| Gene | sgRNA sequences |
| --- | --- |
| *sgSLC15A4*_F | CACCGGGAGCGATCCTGTCGTTAGG |
| *sgSLC15A4*_R | AAACCCTAACGACAGGATCGCTCCC |
| *sgLAMTOR1*_F | CACCGCTGCTACAGCAGCGAGAACG |
| *sgLAMTOR1*_R | AAACCGTTCTCGCTGCTGTAGCAGC |
| *sgTASL*_ F | CACCGGTAGAAATGGAATCCTCCAT |
| *sgTASL*_ R | AAACATGGAGGATTCCATTTCTACC |

**Table S2. List of site-directed mutagenesis primers used in this study.**

| Δ94-161_F | TGGCTGTGGCGGCCGCAGGAGGTGGA |
| --- | --- |
| Δ94-161_R | CGGCCGCCACAGCCAAGCGGGTGCT |
| Δ40-50_F | AGCCCAACCAGGCCCTGCTCTCTTCC |
| Δ40-50_R | GGGCCTGGTTGGGCTCGGCTCCATTG |
| Δ40-65_F | AGCCCAACATTGATGTGTCTGCTGCAGA |
| Δ40-65_R | CATCAATGTTGGGCTCGGCTCCATTG |
| Δ40-80_F | AGCCCAACTACATGGACCGTGCCAGGC |
| Δ40-80_R | CCATGTAGTTGGGCTCGGCTCCATTG |
| Δ40-95_F | AGCCCAACAGCAGCAGCCTGACCCAT |
| Δ40-95 _R | TGCTGCTGTTGGGCTCGGCTCCATTG |
| E50A-F | CACTGATGCCCAGGCCCTGCTCTCTTCC |
| E50A-R | GCCTGGGCATCAGTGCGAGCGGAAGG |
| Q51A-F | TGATGAGGCCGCCCTGCTCTCTTCCATCC |
| Q51A-R | AGGGCGGCCTCATCAGTGCGAGCGGAAG |
| L53/54A-F | AGGCCGCCGCCTCTTCCATCCTTGCCAAGACAG |
| L53/54A-R | AAGAGGCGGCGGCCTGCTCATCAGTGCG |
| S55/56A-F | TGCTCGCCGCCATCCTTGCCAAGACAGCCAGC |
| S55/56A-R | GGATGGCGGCGAGCAGGGCCTGCTCATCAGTGC |
| I57A-F | CTCTTCCGCCCTTGCCAAGACAGCCAGCAAC |
| I57A-R | GCAAGGGCGGAAGAGAGCAGGGCCTG |
| L58A-F | TTCCATCGCCGCCAAGACAGCCAGCAAC |
| L58A-R | TTGGCGGCGATGGAAGAGAGCAGGGCCTG |
| K60A-F | CCTTGCCGCCACAGCCAGCAACATCATTGATG |
| K60A-R | GCTGTGGCGGCAAGGATGGAAGAGAGCAG |
| T61A-F | TGCCAAGGCCGCCAGCAACATCATTGATGTGTCTG |
| T61A-R | CTGGCGGCCTTGGCAAGGATGGAAGAGAGC |
| S63A-F | GACAGCCGCCAACATCATTGATGTGTCTGCTGCAG |
| S63A-R | ATGTTGGCGGCTGTCTTGGCAAGGATGG |
| N64A-F | AGCCAGCGCCATCATTGATGTGTCTGCTGCAG |
| N64A-R | ATGATGGCGCTGGCTGTCTTGGCAAGG |
| SLC15A4_E465K_F | GCATCAGCAAGATCTTCGCCTCCATCGCC |
| SLC15A4_E465K_R | AGATCTTGCTGATGCCGATCAGCAG |
| EcoRI-hygro-F | CGGGGTACCATGAAGAAGCCCGAACTCA |
| KpnI-hygro-R | GGCGAATTCTTAAACTCGACCTACCTCCTTAGCG |

**Table S3. List of qPCR primers used in this study.**

| Gene | Forward primer | Reverse primer |
| --- | --- | --- |
| *IL6* | 5’-AGACAGCCACTCACCTCTTCAG | 5’-TTCTGCCAGTGCCTCTTTGCTG-3’ |
| *TNFA* | 5’-CTCTTCTGCCTGCTGCACTTTG | 5’-ATGGGCTACAGGCTTGTCACTC-3’ |
| *IFNB1* | 5’-GTCACTGTGCCTGGACCATAG-3’ | 5’-GTTTCGGAGGTAACCTGTAAGTC-3’ |
| *CXCL10* | 5’-GGTGAGAAGAGATGTCTGAATCC-3’ | 5’-GTCCATCCTTGGAAGCACTGCA-3’ |
| *STAT1* | 5’-ATGGCAGTCTGGCGGCTGAATT-3’ | 5’-CCAAACCAGGCTGGCACAATTG-3’ |
| *ISG15* | 5’-CTCTGAGCATCCTGGTGAGGAA-3’ | 5’-AAGGTCAGCCAGAACAGGTCGT-3’ |
| *GAPDH* | 5’-GTCTCCTCTGACTTCAACAGCG-3’ | 5’-ACCACCCTGTTGCTGTAGCCAA-3’ |

**Table S4. Key reagents.**

| Reagent | Catalog Number | Supplier |
| --- | --- | --- |
| Antibodies |  |  |
| SLC15A4 polyclonal antibody | This study | Genscript |
| HA-Tag Antibody (F-7) | sc-7392 | Santa Cruz Biotechnology |
| HA-Tag (C29F4) | 3724 | Cell Signaling Technology |
| Phospho-p70 S6 Kinase (Thr389) (108D2) | 9234 | Cell Signaling Technology |
| p70 S6 Kinase (E8K6T) | 34475 | Cell Signaling Technology |
| Phospho-STAT1 (Tyr701) (58D6) | 9167 | Cell Signaling Technology |
| STAT1 polyclonal | 10144-2-AP | Proteintech Group |
| IRF5 polyclonal | 10547-1-AP | Proteintech Group |
| IRF7 (242613C2) | 85072-5-RR | Proteintech Group |
| TASL (CXorf21) polyclonal | 30244-1-AP | Proteintech Group |
| LAMTOR1 Rabbit pAb | A21557 | Abclonal Technology |
| LAMTOR2/ROBLD3 (D7C10) | 8145 | Cell Signaling Technology |
| MAPKSP1 (LAMTOR3) Polyclonal | 11937-1-AP | Proteintech Group |
| LAMTOR4/C7orf59 (D4P6O) | 13140 | Cell Signaling Technology |
| Anti-TLR7 [EPR2088(2)] | ab124928 | Abcam |
| Anti-TLR9 [EPR21735] | ab211012 | Abcam |
| Toll-like Receptor 9 (D9M9H) | 13674 | Cell Signaling Technology |
| Purified anti-human CD289 (TLR9) (S16013D)  (for ICFC & FC) | 394802 | Biolegend |
| RRAGC polyclonal | 26989-1-AP | Proteintech Group |
| RRAGB polyclonal | 13023-1-AP | Proteintech Group |
| ATP6V0D1 polyclonal | 18274-1-AP | Proteintech Group |
| RAB5A polyclonal | 11947-1-AP | Proteintech Group |
| RAB7A polyclonal | 55469-1-AP | Proteintech Group |
| Raptor polyclonal | 20984-1-AP | Proteintech Group |
| Mouse anti Myc-Tag mAb (for ICFC) | AE010 | Abclonal Technology |
| Rabbit anti Myc-Tag pAb | AE009 | Abclonal Technology |
| Purified anti-human CD107b (LAMP-2) (H4B4) | 354301 | Biolegend |
| mTOR (7C10) | 2983 | Cell Signaling Technology |
| anti-human CD107a (LAMP-1) (H4A3) |  | Biolegend |
| CoraLite® Plus 750-conjugated GAPDH (1E6D9) | CL750-60004 | Proteintech Group |
| CoraLite® Plus 750-conjugated Alpha Tubulin (1E4C11) | CL750-66031 | Proteintech Group |
| CoraLite® Plus 750-conjugated Beta Actin (2D4H5) | CL750-66009 | Proteintech Group |
| CoraLite® Plus 750-conjugated Lamin B1 (3C10G12) | CL750-66095 | Proteintech Group |
| PE anti-RPS6 Phospho (Ser235/Ser236) (A17020B) | 608603 | Biolegend |
| APC anti-human CD107a (LAMP-1) (H4A3) | 328619 | Biolegend |
| Alexa Fluor® 488 anti-human CD107a (LAMP-1) (H4A3) | 328609 | Biolegend |
| Alexa Fluor® 647 anti-rat IgG2a (MRG2a-83) | 407511 | Biolegend |
| Alexa Fluor® 488 Donkey anti-rabbit IgG (minimal x-reactivity) (Poly4064) | 406416 | Biolegend |
| Alexa Fluor® 555 Goat anti-mouse IgG (minimal x-reactivity) (Poly4053) | 405324 | Biolegend |
| Purified Rabbit Polyclonal Isotype Ctrl Antibody | 910801 | Biolegend |
| Purified Mouse IgG1, κ isotype Ctrl (MOPC-21) | 400101 | Biolegend |
| Chemicals, peptides, and recombinant proteins |  |  |
| Human IFN-beta Recombinant Protein | 300-02BC-5UG | PeproTech |
| Resiquimod (R848) | HY-13740 | MedChemExpress |
| Agatolimod sodium (ODN2006, CpG-B) | HY-150218 | MedChemExpress |
| ODN 2216 sodium (CpG-A) | HY-150741C | MedChemExpress |
| Poly(I:C) (HMW) | tlrl-pic | Invivogen |
| Dulbecco's Modified Eagle Medium (DMEM) | 11995073 | Thermo Fisher Scientific |
| RPMI 1640 Medium | 11875093 | Thermo Fisher Scientific |
| Trypsin-EDTA (0.05%), phenol red | 25300054 | Thermo Fisher Scientific |
| FBS | FB-01 | Omega Scientific |
| Penicillin-Streptomycin | 15140122 | Thermo Fisher Scientific |
| L-Glutamine (200 mM) | 25030081 | Thermo Fisher Scientific |
| Reagent |  |  |
| Protein A/G Magnetic Beads | HY-K0202 | MedChemExpress |
| Anti-HA Magnetic Beads | HY-K0201 | MedChemExpress |
| RIPA Lysis buffer | 89900 | Thermo Fisher Scientific |
| IGEPAL CA-630 for molecular biology | 9002-93-1 | Sigma-Aldrich |
| Poly-ʟ-Lysine Hydrobromide | P9155-5MG | Sigma-Aldrich |
| Thickness No. 1.5H (tol. ± 5 μm), 24 x 60 mm | 71861-055 | Electron Microscopy Sciences |
| mPAGE® 4X LDS Sample Buffer | MPSB-10ML | Sigma-Aldrich |
| SuperBlock™ Blocking Buffer (T20) | 37536 | Thermo Fisher Scientific |
| Albumin, Bovine Fraction V [BSA], 100 Grams | A30075-100.0 | RPI science |
| Trypsin/Lys-C Mix, Mass Spec Grade | V5071 | Promega |
| Tris(2-carboxyethyl)phosphine hydrochloride | C-1818 | Biosynth |
| Iodoacetamide | I6125 | Sigma-Aldrich |
| Phosphatase Inhibitor Cocktail II | HY-K0022 | MedChemExpress |
| Halt Protease Inhibitor | 78438 | Thermo Fisher Scientific |
| PEI MAX | 24765-1 | Polysciences Inc. |
| CHAPS, Ultra Pure | #22024-10g | Biotium |
| True-Phos™ Perm Buffer | 425401 | Biolegend |
| Critical commercial assays |  |  |
| LumiKin Xpress hIFN-β 2.0 | luex-hifnbv2 | InvivoGen |
| ELISA MAX™ Deluxe Set Human IL-6 | 430504 | Biolegend |
| ELISA MAX™ Deluxe Set Human TNF-α | 430204 | Biolegend |
| BD Cytofix/Cytoperm™ Fixation/Permeabilization Kit | 554714 | BD Bioscience |
| mPAGE®TurboMix Bis-Tris Gel Casting Kit | TMKIT | Sigma-Aldrich |
| TMT10plex™ Isobaric Label Reagents and Kits | 90110 | Thermo Fisher Scientific |
| PowerUp™ SYBR™ Green Master Mix for qPCR | A25742 | Thermo Fisher Scientific |
| GeneJET RNA Purification Kit | K0731 | Thermo Fisher Scientific |
| Zyppy Plasmid Miniprep Kit | D4019 | Zymo Research |
| PrimeScript RT Reagent Kit (Perfect Real Time) | RR037A | Takara Bio |
| Software and algorithms |  |  |
| Proteome Discoverer | Opton | Thermo Scientific |
| GraphPad v10 |  | Dotmatics |
| FlowJo™ v10 |  | FlowJo |
| ImageJ 1.48q |  | doi:10.1038/nmeth.2089 |
| Zen2011 |  | Zeiss |

**Dataset S1 (separate file).** Mass spectrometry-based proteomics analysis dataset.
